# Endoglin, a novel biomarker and therapeutical target to prevent malignant peripheral nerve sheath tumor growth and metastasis

**DOI:** 10.1101/2022.08.12.503580

**Authors:** Teresa González-Muñoz, Angela Di Giannatale, Susana García-Silva, Vanesa Santos, Sara Sánchez-Redondo, Claudia Savini, Osvaldo Graña-Castro, Carmen Blanco-Aparicio, David J Pisapia, Jose L Rodríguez-Peralto, Cleofé Romagosa Pérez-Portabella, Rita Alaggio, Maria Serena Benassi, Laura Pazzaglia, Katia Scotlandi, Nancy Ratner, Kaleb Yohay, Charles P. Theuer, Héctor Peinado

**Author notes:** Corresponding Author, Spanish National Cancer Research Centre (CNIO), Melchor Fernández Almagro, 3, 28029 Madrid, Spain.

## Abstract

**Purpose:** Malignant peripheral nerve sheath tumors (MPNSTs) are highly aggressive softtissue sarcomas that lack effective treatments, underscoring the urgent need to uncover novel mediators of MPNST pathogenesis that may serve as potential therapeutic targets. Tumor angiogenesis is considered a critical event in MPNST transformation and progression. Here, we have investigated whether endoglin (ENG), a TGF-β coreceptor with a crucial role in angiogenesis, could be a novel therapeutic target in MPNSTs.

**Experimental Design:** ENG expression was evaluated in human peripheral nerve sheath tumor tissues and plasma samples. Effects of tumor cell-specific ENG expression on gene expression, signaling pathways and *in vivo* MPNST growth and metastasis were investigated. The efficacy of ENG targeting in monotherapy or in combination with MEK inhibition was analyzed in xenograft models.

**Results:** ENG expression was found to be upregulated in both human MPNST tumor tissues and plasma circulating small extracellular vesicles. We demonstrated that ENG modulates Smad1/5 and MAPK/ERK pathway activation and pro-angiogenic and pro-metastatic gene expression in MPNST cells and plays an active role in tumor growth and metastasis *in vivo*. Targeting with ENG-neutralizing antibodies (TRC105/M1043) decreased MPNST tumor growth and metastasis in xenograft models by reducing tumor cell proliferation and angiogenesis. Importantly, combination of anti-ENG therapy with MEK inhibition reduced more effectively tumor cell growth and angiogenesis.

**Conclusions:** Our data unveil a tumor-promoting function of ENG in MPNSTs and support the combined use of anti-ENG antibodies and MEK inhibitors as a novel potential combination to control MPNSTs growth and metastasis.

**Statement of translational relevance:** Malignant peripheral nerve sheath tumors (MPNSTs) present a poor clinical outcome due to tumor aggressiveness and the absence of effective treatments that underline the need for identifying novel therapeutic approaches. Here, we show that ENG is upregulated in human MPNSTs and we uncover an important role for this coreceptor in MPNST. ENG-neutralizing antibodies (TRC105/M1043) decreased MPNST tumor growth and metastasis in xenograft models, supporting a novel use of anti-ENG therapies for MPNST treatment. Currently, MEK inhibitors are used in the clinic to shrink peripheral nerve sheath tumors (e.g. plexiform neurofibromas). Importantly, we demonstrate that combination of anti ENG antibodies with MEK inhibitors efficiently blocked MPNST growth and metastasis. Our findings provide a rationale for combining anti-ENG and MEK inhibitors as a new strategy for MPNST management.

## INTRODUCTION

MPNSTs are highly aggressive soft-tissue sarcomas (STS) derived from cells of the Schwann cell lineage (1). About 40% to 50% of these tumors arise in the context of the genetic disorder neurofibromatosis type 1 (NF1) (2). Indeed, NF1 patients are predisposed to develop benign peripheral nerve sheath tumors (PNSTs), including dermal and plexiform neurofibromas (PNs) (3). However, plexiform neurofibromas (PNs) can undergo malignant transformation, progressing to pre-malignant lesions called atypical neurofibromatous neoplasms of uncertain biological potential (ANNUBPs) and ultimately MPNSTs, a malignant soft tissue sarcoma with a high mortality rate (3). Another 40% to 47% of MPNSTs are sporadic, and the remaining 10% to 13% occur at sites of prior radiotherapy (2). Irrespective of their origin, these tumors have a strong tendency to relapse and metastasize, with an estimated 5-year survival rates of 15% to 50% (4). Despite this poor prognosis, there is a significant lack of diagnostic biomarkers for early detection of this disease. Regarding the treatment, complete surgical resection, which is the only known effective therapy, is often not feasible due to tumor size, location and/or the presence of metastases. MPNSTs are relatively refractory to conventional radiation and chemotherapy (5) and targeted therapies have shown limited clinical efficacy in MPNSTs (6). Therefore, MPNST patients face a significant unmet therapeutic need, highlighting the urgent demand to uncover novel mediators of pathogenesis and new therapeutic targets.

A common event in the pathogenesis of MPNSTs is the biallelic loss of *NF1* gene that encodes the RAS-GTPase activating protein (GAP) neurofibromin, resulting in the hyperactivation of RAS and its downstream signaling pathways (e.g. RAS/MEK/ERK pathway) (3). In fact, MEK albeit transiently tumor growth in both NF1-associated and sporadic MPNST mouse models (6), supporting the relevance of the RAS/MEK/ERK pathway in MPNST growth. Notably, MEK inhibitors (MEKi) represent the most effective targeted therapy against PNs, recognized precursors of MPNSTs, and they are being tested in clinical trials for MPNSTs (6). In addition to tumor-cell intrinsic mechanisms, alterations in the tumor microenvironment (TME) are now recognized as critical elements influencing MPNST development (7). Particularly, tumor angiogenesis plays a crucial role in the transformation and progression of MPNSTs (8). In fact, increased VEGF expression and vascularization were correlated with tumor malignancy and poor prognosis in MPNST patients (9). A recent phase II trial with the antiangiogenic agent pazopanib demonstrated better outcomes than those achieved with any of the targeted therapies previously tested in MPNST patients, with a clinical benefit rate (CBR) at 12 weeks of 50% (10). Therefore, finding novel players of MPNST angiogenesis should open up additional therapeutic opportunities.

The TGF-β coreceptor endoglin (ENG) is highly expressed in activated endothelial cells (ECs) and it plays a crucial role in angiogenesis (11). Upon stimulation with ligands (e.g. TGF-β and BMP-9), ENG promotes phosphorylation of downstream Smad1 protein, stimulating EC proliferation and migration (11). Notably, ENG expression has been shown to be upregulated in the vascular endothelium of many solid tumor types, being associated with poor prognosis and metastatic disease (12). ENG is also present at high levels in tumor cells of diverse malignancies, including sarcoma (12,13). In fact, ENG promotes the aggressiveness and tumorigenesis of different sarcoma cell lines, and its overexpression predicts worse outcomes in patients with selective sarcoma subtypes (13). However, the role of ENG in MPNST aggressiveness and its potential as a therapeutic target have not been evaluated.

Here we discovered that ENG is highly expressed in both ECs and tumor cells of human MPNSTs, and its expression correlates with advanced tumor stages (local recurrence and distant metastasis). Moreover, ENG levels were elevated in plasma-circulating small extracellular vesicles (sEVs) from MPNST patients. Mechanistically, we found that ENG expression in MPNST cells regulates Smad1/5 and MAPK/ERK pathway activation driving proangiogenic and pro-metastatic gene expression *in vitro*, promoting tumor growth and metastasis *in vivo* as well. Importantly, we reveal that targeting ENG with neutralizing antibodies reduces MPNST tumor growth in xenograft models, by impairing tumor cell proliferation, angiogenesis and ultimately metastasis. Furthermore, the combination of anti-ENG antibodies and the MEKi PD-0325901/Mirdametinib efficiently inhibited both primary tumor growth and metastasis formation. Altogether, our data unveil an important role for ENG in MPNST, promoting tumor cell growth, angiogenesis and metastasis. Notably, our findings suggest that the combination of anti-ENG and MEKi could be a novel approach for the treatment of these tumors.

## MATERIALS AND METHODS

### Patient samples

Human studies were performed according to protocols approved by the Ethical Committee of the Instituto de Salud Carlos III (ISCIII, CEI PI13_2015v2). Paraffin-embedded PNST tissue samples were collected from four independent patient cohorts at Weill Cornell Medical College (New York, USA), Hospital 12 de Octubre (Madrid, Spain), Bambino Gesù Children’s Hospital (Rome, Italy) and Hospital Vall d’Hebron (Barcelona, Spain). The complete collection comprised 58 benign neurofibromas, 9 ANNUBPs and 38 MPNSTs (24 NF1-associated and 14 sporadic). The tissue microarray (TMA) containing 20 tissue cores retrieved from human primary and local recurrent MPNSTs and lung metastases was obtained from the Hospital Virgen del Rocío (Seville, Spain). Human peripheral blood samples were taken from 32 control healthy subjects and patients with PNs (n=32) and MPNSTs (n=11) at Weill Cornell Medical College, Banco Nacional de ADN Carlos III (Salamanca, Spain) and Instituto Ortopedico Rizzoli (Bologna, Italy), all histologically confirmed. All individuals provided informed consent. Whole-blood was collected in EDTA tubes, and plasma was isolated by centrifugation at 1,100g for 10min.

### sEV isolation and ELISA analysis of soluble and EV-secreted ENG in plasma sample

Human plasma samples were centrifuged at 3,000g for 20min. The supernatant was further centrifuged at 12,000g for 20min. sEVs were subsequently harvested by centrifugation at 100,000g for 70min. The supernatant (EV-depleted plasma) was kept to quantify soluble ENG concentration. The sEV pellet was washed with PBS and collected by another ultracentrifugation at 100,000g for 70min. All centrifugations were performed at 10ºC using a Beckman Optima X100 centrifuge with a Beckman 50.4 Ti rotor. sEVs were resuspended in PBS and the protein content was measured by Pierce bicinchoninic acid (BCA) assay (Thermo Fisher). Particle content was determined using Nanosight Nanoparticle tracking analysis (NTA, Malvern). ENG levels were measured in both sEV-enriched and EV-depleted plasma fractions by ELISA (Abcam, ab100507), following the manufacturer’s instructions. For the quantification of sEV-secreted ENG, 5 μg of total sEVs were resuspended in 100 μl of Assay Diluent provided with the kit.

### Cell culture

The human NF1-associated MPNST cell lines ST88-14, 90-8, NMS-2, S462 and the sporadic MPNST cell line STS26T were generously provided by Dr. Eduard Serra (Institute for Health Science Research Germans Trias i Pujol, Barcelona, Spain). The SNF96.2, SNF02.2 and SNF94.3 cell lines (NF1-derived MPNST) were purchased from ATCC. All these cells were cultured in high-glucose DMEM supplemented with 10% FBS, 1 mM sodium pyruvate and 20 μg/ml gentamicin (all from Sigma) at 37 ºC in a humidified atmosphere with 5% CO_2_. All cell lines were routinely tested for mycoplasma contamination.

### Gene silencing by lentiviral transduction

For shRNA-mediated knockdown of *ENG*, lentiviral particles encoding shRNA designed to silence human *ENG* (shENG) or a non-targeting control scrambled sequence (shScramble) were purchased from Santa Cruz Biotechnology. MPNST cells were transduced with the lentiviruses at a low MOI. After 48 hours, infected cells were selected by incubation with 1 μg/ml puromycin (Sigma).

### Cell signaling assays

2×10^5^ MPNST cells were seeded on 6-well plates. Upon 90% confluency, cells were serum-starved overnight in the presence of the anti-human ENG monoclonal antibody (mAb) TRC105 (TRACON Pharmaceuticals), the MEKi PD-0325901 (CNIO Experimental Therapeutics Unit, ETP), the combination of both drugs or IgG control (Jackson ImmunoResearch). TRC105 and IgG were resuspended in PBS and used at a final concentration of 50 μg/ml, and PD-0325901 was reconstituted in DMSO and used at a final concentration of 10 nM for STS26T cells or 1 nM for ST88-14 cells to analyze the synergy between the two compounds. Next day, cells were stimulated with 50 ng/ml recombinant human BMP-9 (R&D Systems) and 150 ng/ml recombinant human VEGF (PeproTech) for one hour. Cells were lysed, protein content was determined, and Western blot analysis was performed (see below).

### Animal studies

All mouse work was approved by the Institutional Ethics Committees of the CNIO (IACUC-006- 2016), the ISCIII (CBA-15_2017) and the Comunidad Autónoma de Madrid (PROEX 168/17). 6-to 8-week-old female athymic nude and NOD scid gamma (NSG) mice were obtained from ENVIGO or CNIO Animal Facility, respectively.

To analyze the effect of *ENG* depletion in MPNST, nude mice were subcutaneously injected with 1×10^6^ shScramble- or shENG-STS26T cells (in a mix of 1:1 serum-free DMEM/Matrigel). Tumor growth was monitored twice weekly by caliper measurement of the two orthogonal large and small external diameters (a, b), and tumor volume was calculated by: a x b2 x π/6. When tumors reached 1.5 cm^3^, mice were sacrificed, and tumors and lymph nodes (LN) were collected and processed for histology.

For anti-ENG and/or MEKi of subcutaneous xenografts, 1×10^6^ STS26T cells or 10×10^6^ ST88-14 cells (in a mix of 1:1 serum-free DMEM/Matrigel) were injected subcutaneously into both flanks of athymic nude or NSG mice, respectively. One-week later, when palpable tumors were present, mice were treated with the anti-ENG mAbs TRC105 and M1043 (TRACON Pharmaceuticals; 10mg/Kg diluted in PBS, intraperitoneal injection twice weekly) or the MEKi PD-0325901 (CNIO-ETP; in order to see synergy between the two compounds, we selected 2mg/Kg concentration dissolved in 0.5% methylcellulose/0.2% Tween-80, oral gavage three times weekly), in monotherapy or combination regimens, as stated in Results. Control groups received thrice-weekly oral administrations of vehicle and/or biweekly intraperitoneal injections of IgG (Jackson Immunoresearch; 10mg/Kg diluted in PBS). Tumor volume was measured as described above. After three weeks of treatment, mice were sacrificed and tumors were excised and processed for histology or RNA extraction. Sentinel LNs were also collected and analyzed for the presence of metastases by hematoxylin and eosin (H&E) staining.

For lung metastasis studies, 1×10^6^ luciferase-expressing STS26T cells were injected via tail vein in nude mice. One-week later, *in vivo* bioluminescence imaging (BLI) was performed using a Xenogen IVIS-200 machine (PerkinElmer), and mice were subsequently randomized into treatment groups (IgG plus vehicle, PD-0325901 alone, PD-0325901+TRC105/M1043) based on equal lung metastatic burden. Dose and treatment schedule were performed as described above. Metastasis was monitored twice weekly by *in vivo* BLI and, after three weeks of treatment, mice were sacrificed and lungs were imaged *ex vivo*.

### RNA extraction, sequencing and data analysis

Total RNA from tissues or cells was extracted using the TRIZOL Reagent (Invitrogen). After recovery of the aqueous phase using chloroform, DNase treatment and further purification were performed using the RNeasy Mini Kit (Qiagen), following the manufacturer’s instructions. RNA sequencing (RNA-seq) was conducted by the CNIO Genomics Unit. Total RNA (1 μg) from samples was used. The average sample RNA Integrity Number was 9.65 (Agilent 2100 Bioanalyzer). cDNA libraries were generated using the Illumina TruSeq RNA Sample Preparation kit and sequenced on an Illumina HiSeq 2500 sequencer. Single-end 50-bp sequenced reads were analyzed with the nextpresso pipeline (14) as follows: sequencing quality was checked with FastQC v.0.10.1. Reads were aligned to the human genome (GRCh37/hg19) with TopHat-2.0.10 using Bowtie v.1.0.0 and SAMtools v.0.1.1.9, allowing two mismatches and 20 multihits. Differential expression was calculated with Cufflinks v.2.2.1 using the human GRCh37/hg19 transcript annotations from https://ccb.jhu.edu/software/tophat/igenomes.shtml. GSEAPreranked was used to perform GSEA of the described gene signatures on a pre-ranked gene list, setting 1,000 gene set permutations. For RNA-seq from MPNST xenografts, an initial filtering was done with Xenome to remove reads coming from mouse. Only those gene sets with FDR *q*-value < 0.25 were considered significant. RNA-seq data generated have been deposited in the Gene Expression Omnibus under the accession number and password (pending).

### Quantitative real time PCR (qRT-PCR)

Total RNA (1 μg) was first reverse-transcribed using the QuantiTect Reverse Transcription kit (Qiagen). 20 ng of total cDNA were then subjected to qRT-PCR using TaqMan Gene Expression Master Mix and pre-designed TaqMan probes (all from Applied Biosystems, Supplementary Table 1). Assays were performed in triplicates on a 7500 Fast Real-Time PCR System (Applied Biosystems). Relative expression was calculated using the ΔΔCt method. *HPRT1* was used as housekeeping gene.

### Immunoblotting

Cells were lysed in RIPA buffer supplemented with protease/phosphatase inhibitors (Roche). Protein concentration was determined by Pierce BCA assay (Thermo Fisher). Equal amounts of protein were resolved by SDS-PAGE and probed with the primary antibodies listed in Supplementary Table 2. Anti-rabbit and anti-mouse IgG-HRP-conjugated antibodies (GE Healthcare) were used as secondary antibodies. Signals were detected using the ECL Western Blotting Substrate kit (GE Healthcare). The intensities of immunoreactive bands were quantified by densitometry using ImageJ software (NIH).

### Histological analyses

Tissues were fixed in formalin and embedded in paraffin, and 3-μm-thick sections were stained with H&E or processed for immunohistochemistry (IHC). Briefly, sections were deparaffinized and antigen retrieval was performed using Tris-EDTA (pH9). Endogenous peroxidase activity was quenched with 3% hydrogen peroxide, and slides were then incubated with the appropriate primary antibodies (Supplementary table 2). An UltraVision ONE Detection System (Thermo Fisher) was used following the manufacturer’s protocols. Sections were counterstained with hematoxylin (Anatech) and mounted with permanent mounting medium. Image analysis was performed using the ZEN software (Zeiss). ENG staining in human tumor tissues was scored by four independent pathologists according to the intensity and extent of expression (0-3).

### Statistical analysis

All analyses were performed using GraphPad Prism software v.9.1.0. Data are presented as mean ± SEM unless otherwise indicated. Specific number of independent experiments and biological replicates are stated in the figure legends. Differences between two groups were calculated using two-tailed unpaired Student’s *t* test or Mann-Whitney test; for multiple groups, one-way ANOVA or Kruskal-Wallis (nonparametric) test were used. Tumor growth curves were analyzed by two-way ANOVA. *P*-values are indicated in each figure for statistically significant comparisons (p<0.05).

## RESULTS

### ENG is overexpressed in human primary and metastatic MPNSTs and plasma circulating sEVs

Based on the data supporting a pro-tumorigenic role of ENG in sarcomas (12,13), we hypothesized that ENG could be also involved in PNST transformation, growth and/or metastasis. We first analyzed ENG expression by IHC in human benign neurofibromas (n=58), pre-malignant lesions (ANNUBPs, n=9) and MPNSTs (n=38) collected from four independent patient cohorts. We found a high expression of ENG in both tumor cells and ECs in MPNSTs compared to ANNUBPs or benign neurofibromas (**Fig. 1A,B**). Next, we examined ENG expression in a human TMA comprising primary and local recurrent MPNSTs and metastases. Analysis of ENG staining in both tumor cells and ECs showed that the expression of this protein increased in local recurrent tumors, being significantly higher in metastatic lesions compared to primary tumors (**Fig. 1C,D**).

**Figure 1.**
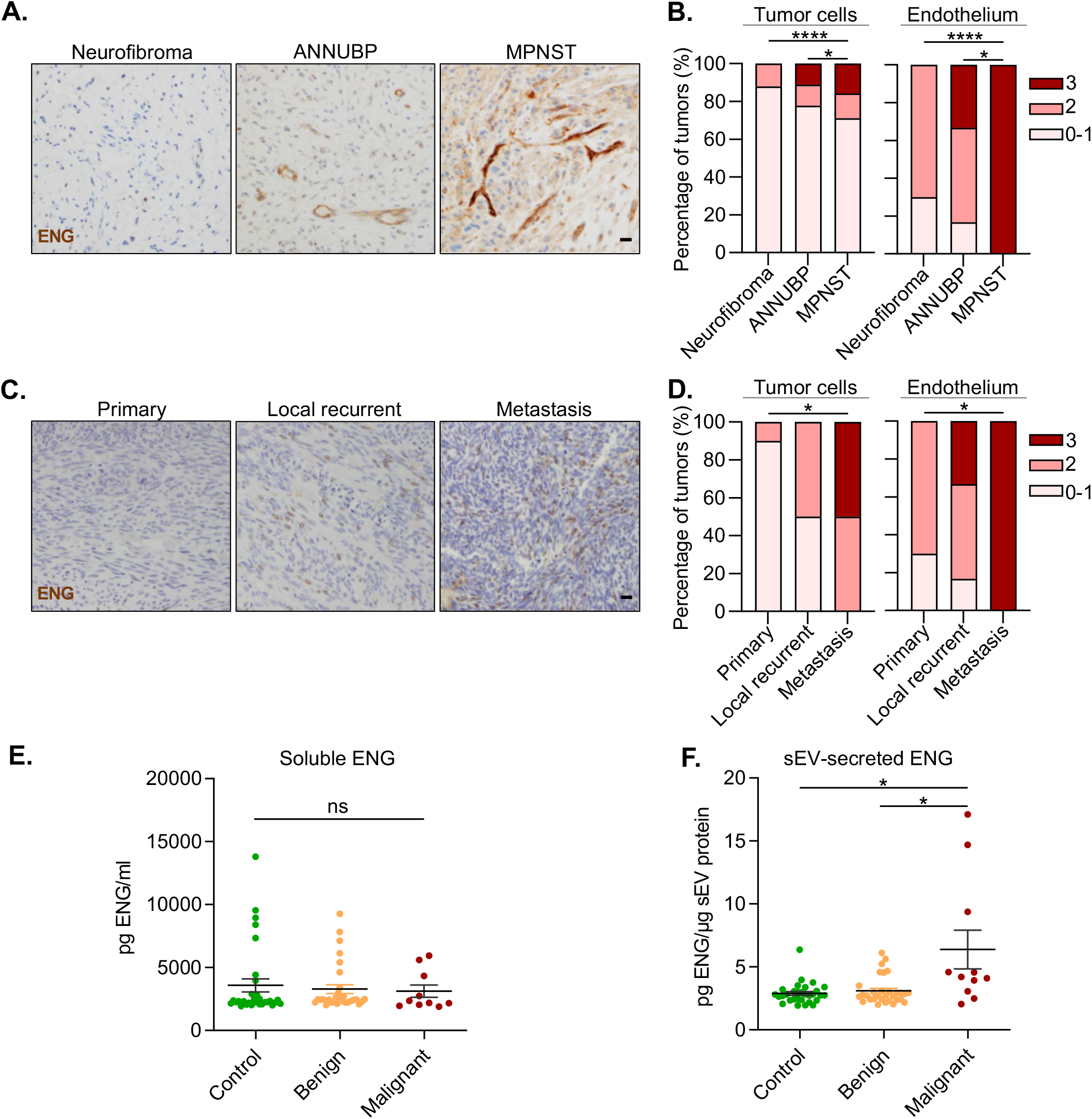
ENG is upregulated in human MPNST tissues and plasma circulating sEVs. **A)** Immunohistochemical staining for ENG in the indicated PNSTs. Scale bar=100μm, and **B)** their distribution (%) according to ENG expression. Score 0-1, low ENG staining; 2, intermediate ENG staining and 3, high ENG staining. Neurofibromas n=58, ANNUBPs n=9, and MPNSTs n=38 from four independent patient cohorts. * P<0.05, **** P<0.0001 by Kruskal-Wallis test. **C)** Representative images of ENG IHC staining in a human TMA comprising primary and locally recurrent MPNSTs and metastases. **D)** Quantification of ENG expression (score 0-3). Primary tumors n=10, local recurrent tumors n=6 and metastases n=2. * P<0.05 by Kruskal-Wallis test. **E**,**F)** Quantification of ENG levels in EV-depleted soluble fraction (**E**) and circulating sEVs (**F**) collected from the plasma of healthy controls (n=23) and patients with benign (PNs, n=33) or malignant (MPNSTs, n=11) PNSTs. sEV-secreted ENG levels were normalized to the total amount of sEV protein. Mean ± s.e.m.; ns, not significant; * P<0.05 by one-way ANOVA.

We evaluated the relevance of circulating ENG in PNST by analyzing the levels of both soluble and EV-shed ENG in the plasma of 44 patients with PNSTs at different stages (benign PNs and MPNSTs) and 23 healthy controls. We separated plasma soluble proteins from sEVs using standard ultracentrifugation methods. sEV number and protein concentration did not differ based on tumor stage (**Supplementary Fig. 1A,B**). Analysis of ENG protein concentration by ELISA showed no significant differences in the expression of soluble ENG between the groups (**Fig. 1E**). However, the levels of ENG in sEVs were significantly increased in MPNST patients compared to patients bearing benign PNs or healthy controls (**Fig. 1F**).

### ENG contributes to the acquisition of malignant traits in MPNST cells

To explore the functional importance of ENG in MPNSTs, we first analyzed its expression in a panel of established human MPNST cell lines. High ENG protein levels were observed in all these cell lines, except for the S462 model (**Fig. 2A**). We then silenced *ENG* expression in STS26T and ST88-14 cells by lentiviral-mediated delivery of shRNAs. STS26T cell line derives from a sporadic MPNST (15) and ST88-14 is a NF1-associated cell line with confirmed loss of heterozygosity at the *NF1* locus (16). We verified that ENG mRNA and protein levels were reduced by 80% in *ENG*-depleted STS26T and ST88-14 cells (shENG) compared to their respective controls (shScramble) (**Fig. 2B,C**).

**Figure 2.**
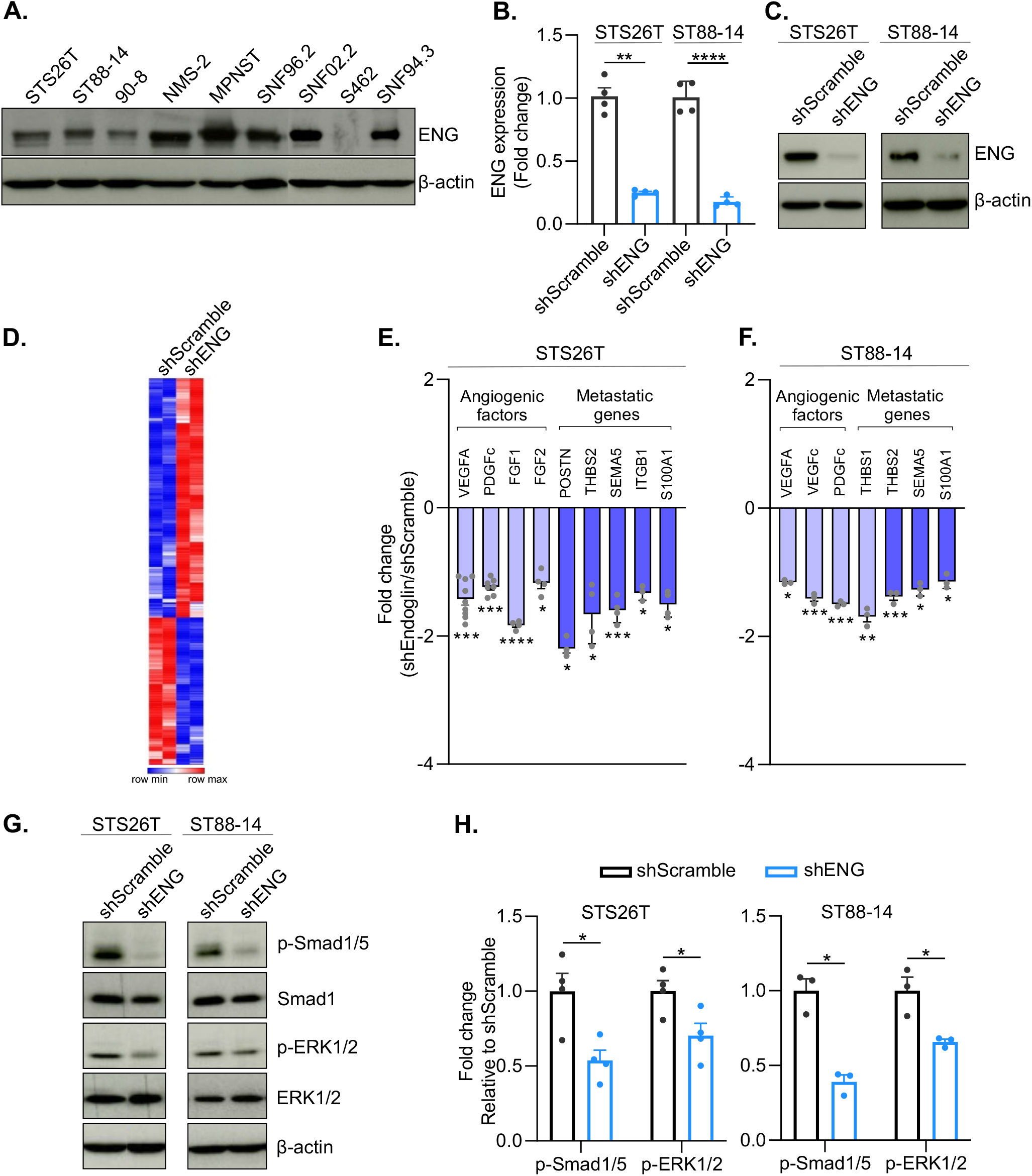
ENG regulates oncogenic pathways in MPNST cells. **A)** Immunoblotting showing ENG protein levels in a panel of human MPNST cell lines. **B)** Quantitative real time PCR analysis of *ENG* expression after shRNA knockdown of *ENG* in the STS26T and ST88-14 cell lines. **C)** Western blot analysis of ENG in STS26T and ST88-14 cells transduced with lentiviral vectors coding for scramble- or ENG-shRNA. **D)** Heat-map representation of the differentially expressed genes in shENG-versus shScramble-STS26T cells (FDR q-value<0.05). **E**,**F)** qRT-PCR validation of the indicated angiogenic and metastatic genes in STS26T (**E**) and ST88-14 (**F**) cells upon *ENG* knockdown. **G**,**H)** Representative Western blot images (**G**) and quantification (**H**) of Smad1/5 and ERK1/2 phosphorylation in shScramble and shENG-STS26T and ST88-14 cells. p-Smad1/5 and p-ERK1/2 levels were normalized to total Smad1 and ERK1/2 levels, respectively. Data in **B**,**E**,**F**,**H** are presented as the fold change compared to shScramble. Mean ± s.e.m. of at least three biological replicates; * P<0.05, ** P<0.01, *** P<0.001, ****P<0.0001 by unpaired t-test.

To examine the gene expression changes related to *ENG* silencing in MPNST cells, we performed RNA-seq after knockdown of *ENG* in STS26T cells compared to shScramble control. *ENG* depletion caused notable changes in the transcriptome of STS26T cells, leading to a significant deregulation of 951 genes (FDR<0.05) (**Fig. 2D** and **Supplementary Table 3**). Gene Set Enrichment Analysis (GSEA) revealed a significant suppression of gene signatures associated with oncogenic traits, including soluble angiogenic factors (e.g. VEGFA, PDGF, PIGF) and metastasis (**Supplementary Fig. 2A,B**). Accordingly, the downregulation of several pro-angiogenic factors (e.g. *VEGF, PDGFc, FGF*) and pro-metastatic genes (e.g. *THBS, POSTN, SEMA5, ITGB1, S100A1*) upon *ENG* depletion was confirmed at the mRNA level in both STS26T and ST88-14 cells (**Fig. 2E, F**).

We next analyzed signaling pathways affected by ENG in MPNST cells. This TGF-β coreceptor is known to modulate TGF-β signaling by promoting the ALK1-Smad1/5 pathway (11). Moreover, ENG can mediate non-canonical, non-Smad pathways such as the MAPK/ERK signaling cascade (17), which plays crucial roles in MPNST malignancy and progression (8). We thus focused on evaluating the importance of ENG for these signaling pathways in MPNST cells. *ENG* silencing significantly reduced the phosphorylation of Smad1/5 and ERK1/2 in both the sporadic cell line STS26T and the NF1-derived cell line ST88-14 (**Fig. 2G,H**).

### ENG promotes MPNST growth and metastasis *in vivo*

We next studied the effect of *ENG* depletion on MPNST *in vivo*. STS26T-shENG- or – shScramble cells were subcutaneously implanted into nude mice. Downregulation of *ENG* significantly reduced xenograft growth (**Fig. 3A**). Moreover, analysis of spontaneous metastasis at end-point (day 32 post-tumor cell injection) showed that the number and size of LN metastases were decreased in mice bearing *ENG*-depleted xenografts compared to those with control tumors (**Fig. 3B**). Western blot and IHC studies confirmed a strong reduction of ENG protein levels in xenografts arising from shENG cells (**Supplementary Fig. 3A,B**). In order to dissect the mechanisms underlying the role of ENG in MPNST *in vivo*, we analyzed the effect of *ENG* knockdown on both tumor cell proliferation and death. We did not found differences in the expression of the apoptotic marker active caspase-3 between shScramble and shENG tumors (**Supplementary Fig. 3C,D**). However, we observed that *ENG* knockdown significantly reduced tumor cell proliferation as denoted by the strong decrease in the number of Ki67^+^ cells (**Fig. 3C,D**). Moreover, *ENG* depletion in tumor cells significantly reduced tumor vascularization in STS26T xenografts, as determined by staining for the EC marker CD31 (**Fig. 3E,F**).

**Figure 3.**
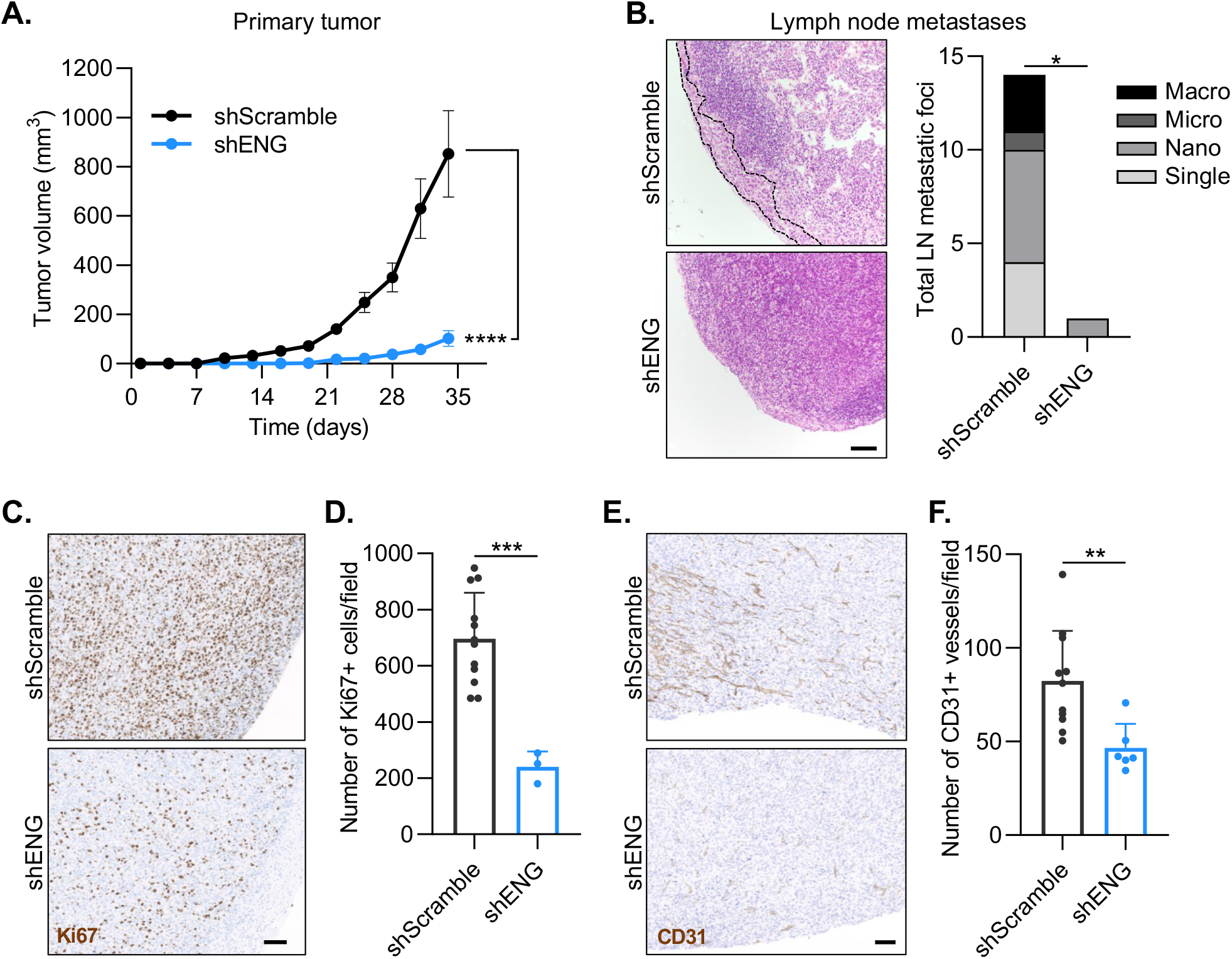
Disrupting *ENG* expression in tumor cells reduces MPNST growth and metastasis. **A)** Analysis of primary tumor growth in shScramble- and shENG-STS26T cells subcutaneously injected into both flanks of athymic nude mice. n=10 mice per group from two independent experiments. Mean ± s.e.m.; **** P<0.0001 by two-way ANOVA. **B)** LN metastasis burden was determined by H&E staining in LN sections at day 32 post-tumor cell injection (end-point). The black points encircle a representative metastasis. Scale bar=100μm. Quantification of the total number of metastatic foci in the LNs (right panel). Lesions were binned into four size categories according to the number of tumor cells: single (1-2 cells), nano (3-10 cells), micro (11-30 cells) and macro (greater than 30 cells). n=10 LNs per group; * P<0.05 by Mann-Whitney test. **C**,**D)** IHC analysis of Ki67 in shScramble- and shENG-STS26T xenografts (Scale bar=100μm) (**C**), and quantification (**D**). **E)** Representative images of CD31 immunohistochemical staining in the indicated tumors. Scale bar=100μm. **F)** Quantification of the number of CD31^+^ vessels. Dots in the graphs in **D**,**F** represent the mean number of positive cells/vessels per field of each tumor (5 fields/tumor). Mean ± s.e.m; ** P<0.01, *** P<0.001 by unpaired t-test.

### Pharmacological targeting of ENG decreases MPNST growth and metastasis

We next ascertained whether ENG could be exploited as potential therapeutic target in MPNSTs. Based on our data showing that ENG is upregulated in tumor cells and also in the vasculature of human MPNSTs, we tested a therapeutic strategy to block ENG on both tumor cells and ECs in MPNST xenografts using a combination of the anti-human and anti-mouse ENG mAbs TRC105 and M1043, respectively (18).

Notably, administration of TRC105/M1043 to nude mice bearing established STS26T xenografts (formed 7 days after tumor cell injection) resulted in a significant decrease in both primary tumor growth and LN metastatic burden (**Fig. 4A,B** and **Supplementary Fig. 4A**). To confirm the efficacy of anti-ENG therapy in an additional MPNST model, we used ST88-14 cells subcutaneously injected into NSG mice. Co-treatment with TRC105/M1043 after tumor establishment (one-week post-tumor cell injection) significantly reduced tumor growth also in this MPNST model and led to tumor regression in 70% of treated mice (**Fig. 4C,D** and **Supplementary Fig. 4B**). These data revealed that ENG-blocking antibodies decreases MPNST growth and metastasis, suggesting that anti-ENG therapies could represent a novel treatment option for this tumor type.

**Figure 4.**
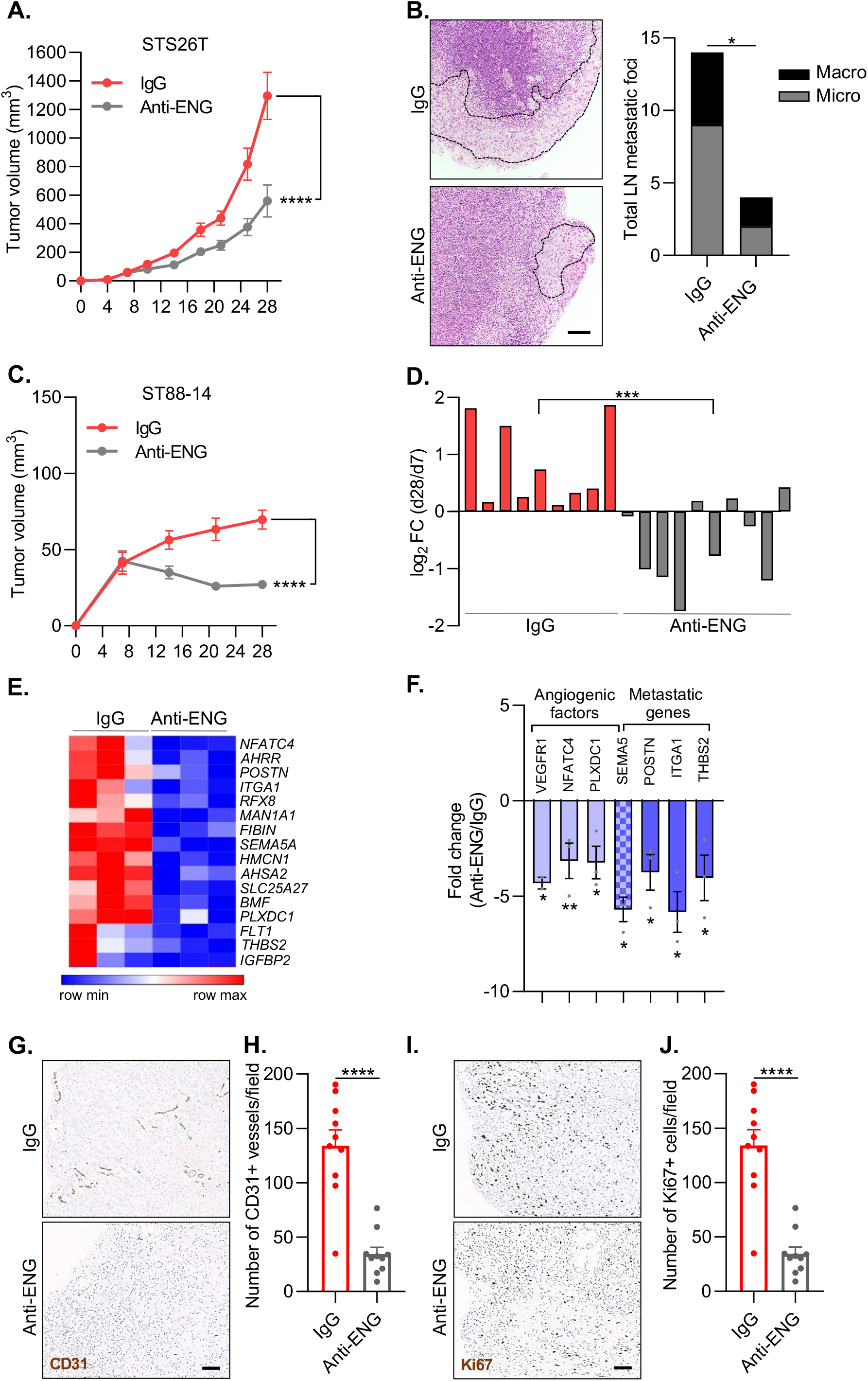
Anti-ENG therapy decreases tumor growth in MPNST xenograft models. **A)** Growth curves of STS26T xenografts treated with TRC105 and M1043 (anti-ENG) or IgG control. n=5 mice per group (10 tumors/treatment). **B)** H&E staining of representative LNs collected from mice bearing STS26T xenografts at the end of the treatment (day 28; left panels). Metastases are highlighted by a discontinuous black line. Scale bar=100μm. Quantification of the total number of metastatic foci (right panel). Lesions were divided into micro-metastasis (11-30 cells) and macro-metastasis (greater than 30 cells). n=10 LNs per group; * P<0.05 by Mann-Whitney test. **C)** Tumor growth of ST88-14 xenografts treated as indicated. n=5 mice per group (10 tumors/treatment). Data in **A**,**C** are presented as mean ± s.e.m.; **** P< 0.0001 by two-way ANOVA. **D)** Waterfall plot showing change in tumor volume (represented as log2 fold change) of individual tumors formed at 28 days after transplant, and after 21 days with pharmacological inhibition. n= 5 mice per group (10 tumors/treatment). ** P< 0.01 by unpaired t-test. **E)** Heat-map showing the significantly downregulated human genes in anti-ENG (TRC105/M1043)-treated STS26T xenografts compared to control IgG-treated tumors. n=3 tumors per group. FDR q-value<0.05. **F)** qRT-PCR validation of the indicated genes obtained from RNA-seq results. Data were normalized to IgG. Mean ± s.e.m. of three biological replicates; * P< 0.05, ** P< 0.01 by unpaired t-test. **G)** Tumor vessel density was assessed by IHC for CD31 in xenografts collected at the completion of treatment. Representative images from each group are shown. Scale bar=100μm. **H)** Quantification of the number of CD31^+^ vessels in tumor sections. **I**,**J)** Representative images of Ki67 immunohistochemical staining in the indicated tumors (**I**) and quantification (**J**). Dots in the graphs in **H, J** represent the mean number of positive cells/vessels per field of each tumor (5 fields/tumor). Mean ± s.e.m.; **** P<0.0001 by unpaired t-test.

To analyze the genes and molecular pathways modulated by ENG targeting, we performed RNA-seq on the STS26T xenografts treated with TRC105/M1043 and compared their expression profiles with those of IgG-treated control tumors. Consistent with our *in vitro* results, several pro-angiogenic and pro-metastatic human genes were significantly downregulated in these tumors upon ENG inhibition (**Fig. 4E**). Accordingly, we found a significant suppression of gene signatures related to angiogenesis and metastasis (**Supplementary Fig. 4C,D**). We validated a subset of pro-angiogenic (e.g. *VEGFR1, NFATC4, PLXDC1*) and pro-metastatic (e.g. *SEMA5, POSTN, ITGA1, THBS2*) genes by qRT-PCR (**Fig. 4F**). Moreover, IHC analysis of CD31 confirmed a significant reduction of tumor vascularization in both STS26T (data not shown) and ST88-14 xenografts treated with TRC105/M1043 (**Fig. 4G,H**). In line with our data using the *ENG* knockdown model, pharmacological targeting of ENG also significantly reduced tumor cell proliferation in both STS26T (data not shown) and ST88-14 tumors (**Fig. 4I,J**), as denoted by the decrease in the number of Ki67^+^ cells.

### Combination therapy with anti-ENG antibodies and the MEKi PD-0325901 efficiently inhibits MPNST growth and metastasis

MEK inhibition is a promising strategy for the treatment of PNSTs, which commonly harbor hyperactivation of the MAPK/ERK signaling pathway (8). Indeed, MEKis have become the first FDA-approved therapy for patients with PNs, recognized precursors of MPNSTs, and they have demonstrated strong albeit transient efficacy in preclinical MPNST models, leading to ongoing clinical trials (e.g. SARC031, (6,19)). We thus set out to evaluate the use of the combination of anti-ENG mAbs with a MEKi.

We first tested the efficacy of this combination treatment in already established primary MPNSTs. STS26T cells were subcutaneously injected into nude mice, and tumors were allowed to develop for one week (∼100 mm^3^ average). Mice were then treated with MEKi PD-0325901 alone, the combination of PD-0325901 with TRC105/M1043 (combo) or IgG plus vehicle (control). Tumor growth was monitored and sentinel LN metastases were assessed at end-point (**Fig. 5A**). Consistent with previously reported data (8), PD-0325901 treatment resulted in decreased MPNST growth and LN metastatic burden (**Fig. 5B-D** and **Supplementary Fig. 5A**). Notably, the combination treatment significantly improved the anti-tumor efficacy of MEKi monotherapy and almost abolished spontaneous LN metastasis (**Fig. 5B-D** and **Supplementary Fig. 5A**). No adverse effects of drug treatment on body weight or skin were observed (**Supplementary Fig. 5B**).

**Figure 5.**
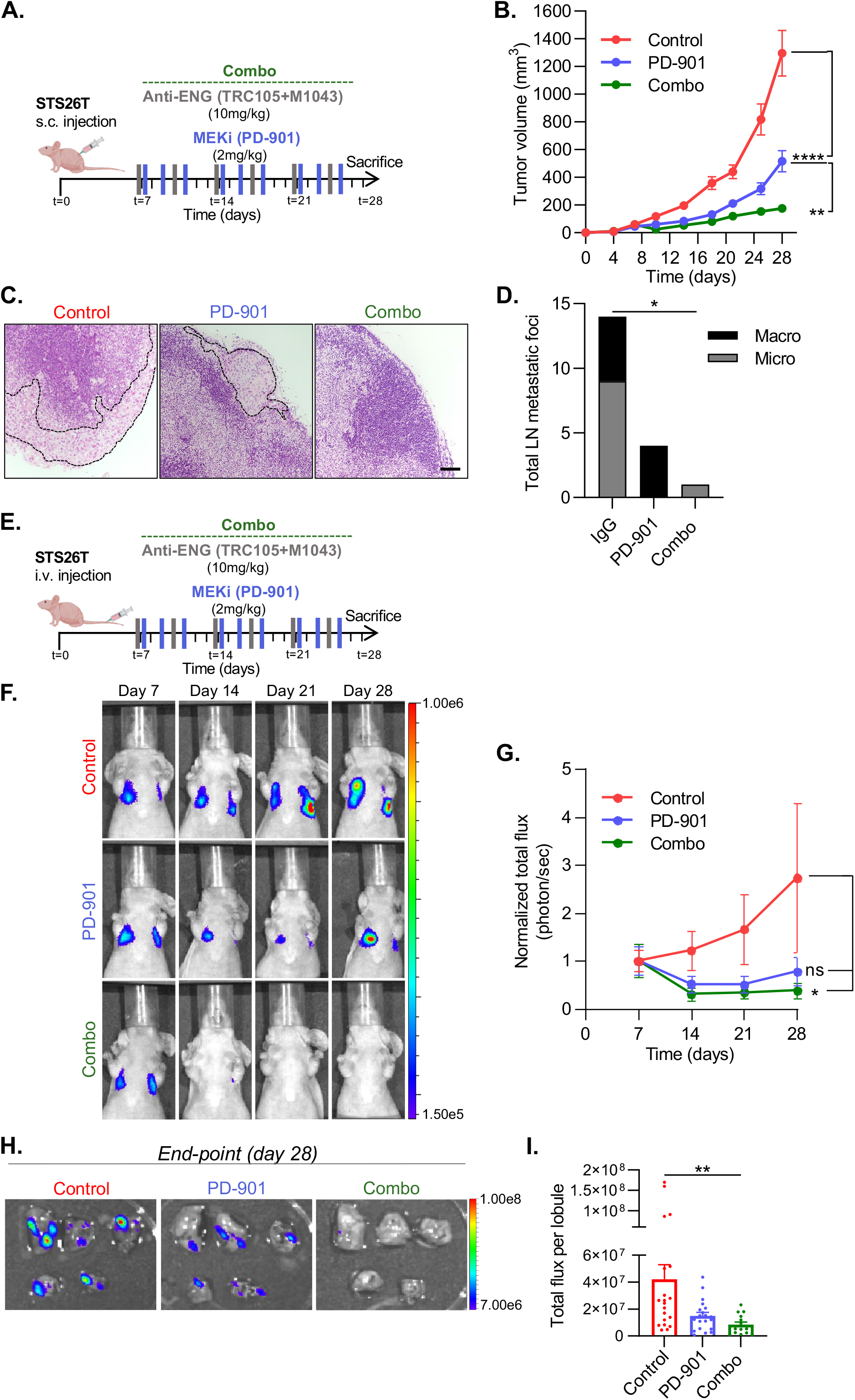
Combination treatment with anti-ENG and anti-MEK therapies inhibits MPNST growth and metastasis. **A)** Schematic of the experiment performed to analyze the efficacy of the anti-ENG/anti-MEK combination therapy in STS26T subcutaneous xenograft model. **B)** STS26T xenograft growth in mice treated with IgG plus vehicle (control), PD-0325901 (PD-901) alone or the combination of PD-901 and the anti-ENG mAbs TRC105 and M1043 (combo). n=5 mice per group (10 tumors/condition). Mean ± s.e.m.; ** P<0.01, ****P<0.0001 by two-way ANOVA. **C)** Analysis of LN metastases by H&E staining at day 28 (end-point). Metastases are highlighted by a discontinuous black line. Scale bar=100μm. **D)** Quantification of the total number of metastatic foci. Lesions were divided into micro-metastasis (11-30 cells) and macro-metastasis (greater than 30 cells). n=10 LNs/group; * P<0.05 by Mann-Whitney test. **E)** Schematic of the experiment performed to evaluate the effect of the anti-ENG/anti-MEK combination therapy on experimental MPNST lung metastasis. **F**,**G)** Representative BLI images (**F**) and quantification (**G**) of experimental metastasis assay follow-up in mice receiving the indicated treatments. Data were normalized to the total flux mean from each group at day 7 (start of treatment). **H)** Assessment of lung metastatic burden by ex vivo BLI at day 28 (end-point), with **I)** quantification of total photon flux per lung lobe. n=4 mice per group. Data are mean ± s.e.m.; * P<0.05 by two-way ANOVA in (**G**) and ** P< 0.01 by one-way ANOVA in (**I**).

Given that up of 50% of MPNST patients develop metastatic disease, normally to the lung (20), we next analyzed the impact of the combination therapy on distant metastasis. Nude mice were injected via tail vein with luciferase-expressing STS26T cells, which exhibit metastatic tropism to the lung (21). One week later, mice with established metastases were distributed into the control and treated groups (PD-0325901 alone, PD-0325901 +TRC105/M1043) based on equal bioluminescent signal (**Supplementary 5C**), and metastatic growth was followed by *In vivo* Imaging System (IVIS, **Fig. 5E**). Although PD-0325901 monotherapy reduced lung metastasis, only the combination treatment with TRC105/M1043 demonstrated a statistically significant decrease in MPNST metastatic outgrowth and lung metastatic burden at end-point (**Fig. 5F-I** and **Supplementary Fig. 5D,E**).

### Targeting of ENG and MEK inhibition cooperatively reduces Smad1/5 and MAPK/ERK pathway activation, MPNST cell proliferation and angiogenesis

To explore the molecular mechanisms responsible for the effectiveness of the anti-ENG and MEKi combination therapy in MPNSTs, we analyzed its effect on ENG-Smad1/5 and MAPK/ERK pathway activation in tumor cells. STS26T and ST88-14 cells were treated with TRC105 alone, PD-0325901 alone, the combination of both or IgG control and subsequently stimulated with BMP-9, the main ENG ligand (22), and VEGF, an important activator of the MAPK/ERK pathway (23). We found that only the TRC105/PD-0325901 combination significantly reduced ENG expression and Smad1/5 phosphorylation in both STS26T and ST88-14 cells (**Fig. 6A,B** and **Supplementary Fig. 6A,B**). Although MEKi treatment increased p-MEK1/2 levels consistent with signaling rebound upon feedback relief (24), it led to diminished phosphorylation of ERK, the direct downstream target of MEK (**Fig. 6A,B** and **Supplementary Fig. 6A,B**). TRC105 alone also produced a significant decrease in p-ERK levels (**Fig. 6A,B** and **Supplementary Fig. 6A,B**). Importantly, dual targeting of ENG and MEK resulted in synergistic inhibition of ERK1/2 phosphorylation in the two MPNST cell lines (**Fig. 6A,B** and **Supplementary Fig. 6A,B**). These data indicate that ENG targeting cooperates with MEK inhibition to block the activation of both ENG-Smad1/5 and MAPK/ERK pathways in MPNST cells.

**Figure 6.**
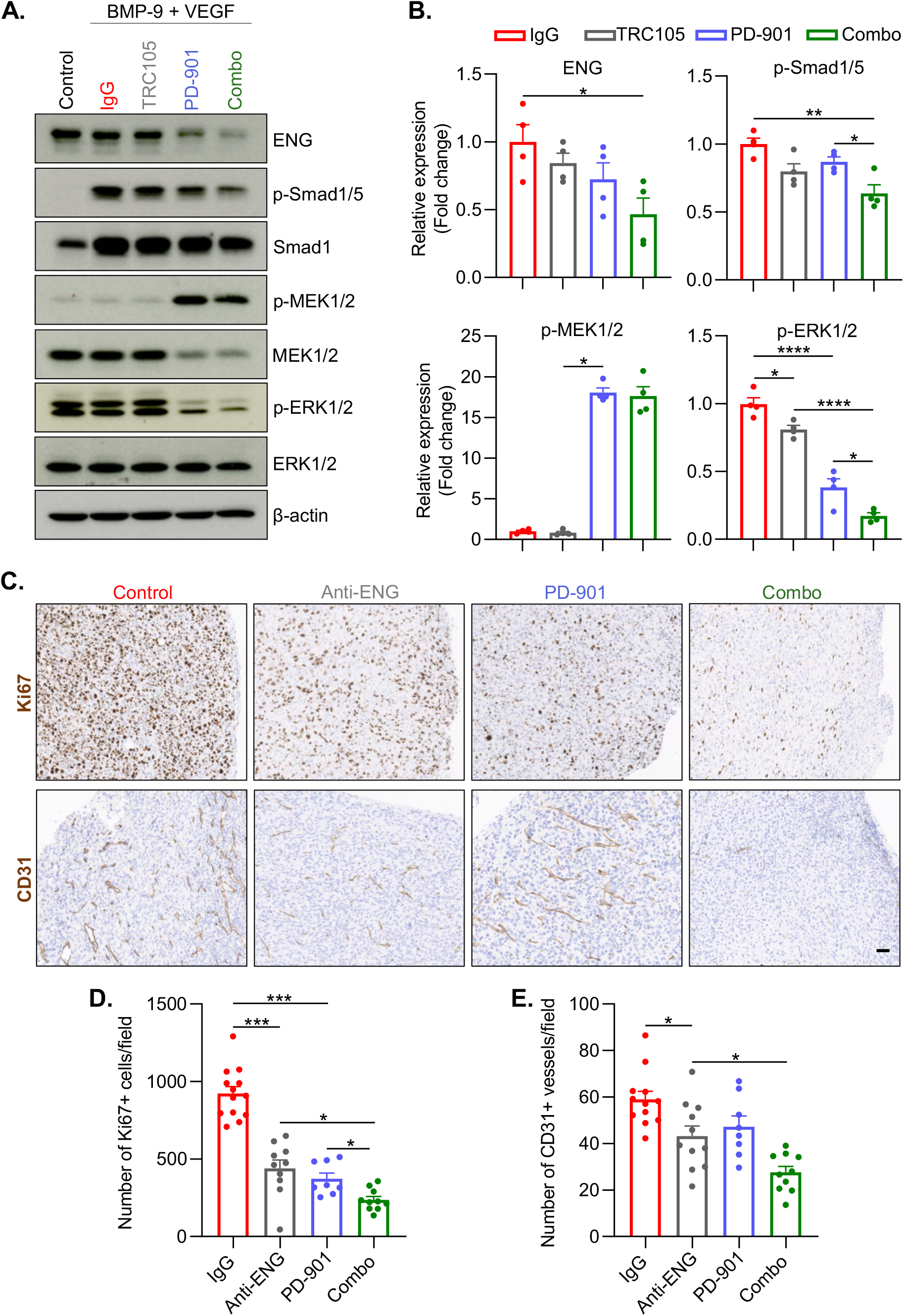
Dual inhibition of ENG and MEK cooperates to reduce Smad1/5 and MAPK/ERK signaling activity, proliferation and angiogenesis. **A, B)** Representative Western blot imagesp>(**A**) and quantification (**B**) of ENG expression and Smad1/5, MEK and ERK activation in STS26T cells pre-treated overnight with a IgG control, the anti-human ENG mAb TRC105 alone, the MEKi PD-0325901 (PD-901) alone or the combination of both drugs (combo) and then stimulated with BMP-9/VEGF for 1 hour. Cells without neither the pre-treatment nor BMP-9/VEGF stimulation were used as a control. Phosphorylated protein levels were normalized to the corresponding total protein levels. Data are presented as the fold change compared to IgG. Mean ± s.e.m. of four biological replicates; * P<0.05, ** P<0.01, **** P<0.0001 by one-way ANOVA. **C)** IHC of Ki67 and CD31 in STS26T xenografts from mice treated with IgG plus vehicle (control), the anti-ENG mAbs TRC105 and M1043 (anti-ENG), PD-901 alone or the combination of TRC105/M1043 with PD-901(combo). **D, E)** Quantification of the number of Ki67^+^ cells (**D**) and CD31^+^ vessels (**E**) in tumor sections (5 fields/tumor). Dots in the graph represent the mean number of positive stained cells/vessels per field of each tumor. Mean ± s.e.m; * P<0.05, *** P<0.001 by one-way ANOVA.

Given the role of these pathways in cancer cell proliferation and angiogenesis (23) together with our data demonstrating that ENG promotes MPNST cell proliferation and vascularization, we next analyzed the effect of dual blockade of ENG and MEK on these biological processes in MPNSTs. IHC analysis of Ki67 showed that both ENG targeting with TRC105/M1043 and MEK inhibition with PD-0325901 significantly reduced tumor cell proliferation in STS26T xenografts (**Fig. 6C,D**). Notably, the combination treatment was significantly more effective than either anti-ENG or MEK inhibitor monotherapy in inhibiting tumor cell proliferation in these tumors (**Fig. 6C,D**). In addition, as indicated above, TRC105/M1043 anti-ENG treatment decreased the number of CD31^+^ vessels in STS26T tumors (**Fig. 6C,E**). These anti-angiogenic effects were significantly enhanced upon combination with PD-0325901 (**Fig. 6C,E**).

## DISCUSSION

The highly aggressive and metastatic behavior of MPNSTs (20) and the absence of effective systemic treatments (5,8) underscore the urgent need to identify novel mediators of pathogenesis that are amenable to therapeutic intervention. In this study, we show that the TGF-β coreceptor ENG is a novel player and a promising therapeutic target in MPNSTs.

ENG expression has been reported in different human sarcomas, correlating with malignancy, aggressiveness or worse survival (25-29). Here we demonstrate that ENG is highly expressed on both tumor cells and ECs in human MPNSTs and that its expression correlates with advanced stages of the disease (local recurrence and distant metastasis), providing evidence for the use of both tumor and endothelial ENG as markers of MPNSTs transformation. Unfortunately, tissue biopsy presents limitations for the management of MPNST patients, mainly due to tumor location and intratumoral heterogeneity (20). Liquid biopsy, which is emerging as a non-invasive method for early cancer detection (30), remains poorly explored in MPNSTs. The use of circulating biomarkers such as circulating-free DNA (cfDNA) (31) or sEVs (this study) may represent an attractive non-invasive approach for early MPNST detection and monitoring, an idea that must be further developed in the field. We found that ENG levels were increased in plasma-circulating sEVs from patients with MPNSTs compared to PN patients or healthy controls. Besides ENG abundance in tumor tissue, high levels of two circulating forms of ENG (soluble and in association with EVs) have been detected in plasma from patients with breast, prostate and colorectal cancer, correlating with disease progression (32-35). To date, only one study has investigated the potential of EV-shed ENG as a circulating biomarker in cancer patients, showing that plasma concentration of ENG^+^ EVs significantly distinguished breast cancer patients from healthy subjects (35). Our data go beyond these previous findings; we propose that an analysis of EV-secreted ENG by liquid biopsy could be useful to detect malignant transformation of PNSTs. Nevertheless, our results on circulating sEVs have to be interpreted with caution, since ENG overexpression was observed only in some MPNST, which could be attributed to the heterogeneity of human disease. Unfortunately, in our series we did not have access to matched tumor and plasma samples or follow-up information. Studies with larger paired tissue/blood specimens, in particular during follow-up and linked clinical data, would be crucial to validate ENG’s potential utility in liquid biopsies.

ENG mainly modulates malignant phenotypes of cancer cells by regulating TGF-β/BMP/Smad signaling but also Smad-independent pathways such as the MAPK/ERK signaling cascade (12), which plays crucial roles in MPNST tumor growth (8). Consistent with other studies in different tumor models (29,36), we found that *ENG* downregulation attenuated Smad1/5 pathway activation in both the sporadic cell line STS26T and the NF1-associated cell line ST88-14. Likewise, *ENG* knockdown led to reduced ERK1/2 phosphorylation in these MPNST cells, which is in agreement with a prior report revealing that *ENG* depletion in uterine leiomyosarcoma cells decreased p-ERK1/2 levels, impairing cell invasion (26). However, it was shown that ENG expression in transformed epidermal cells inhibited the MAPK/ERK pathway, thus suppressing H-RAS-mediated oncogenic transformation (37). Therefore, the precise function of ENG in regulating MAPK/ERK signaling seems to depend on tumor cell type, sarcoma versus carcinoma, which appears to be associated with pro-tumoral or anti-tumoral effects, respectively. Our data, together with previous results proposing that a crosstalk between Smad1/5 and MAPK/ERK signaling pathways promotes MPNST cell aggressiveness (38), suggest a role for ENG in favoring the communication between these pathways, thus potentially contributing to malignant behavior of MPNST cells.

Indeed, we demonstrate that ENG expression in MPNST cells favors tumor growth and metastasis *in vivo*, as *ENG* knockdown resulted in reduced tumor cell proliferation and impaired tumor growth and metastasis. Accordingly, previous studies showed that *ENG* depletion decreased the metastatic abilities of different sarcoma cell lines (25,26,29,39) and inhibited *in vivo* progression of Ewing sarcoma cells (29), indicating a pro-tumoral action of ENG in sarcoma cells. However, the role of ENG in carcinoma cells is context-dependent, in some cases (e.g. melanoma, pancreatic, renal and ovarian cancer) promoting tumor progression and aggressiveness, whereas in other cases (e.g. spindle cell carcinoma, esophageal, lung and breast cancer) it has been associated with tumor suppression (12). These findings point to specific effects of ENG on tumorigenesis depending on the type/origin of tumor cells (mesenchymal versus epithelial), which could be associated with differences in ENG expression levels and modulation of Smad and non-Smad pathways. Interestingly, although endothelial ENG has been widely reported to promote vascularization in different cancer mouse models (18,40-42), our study shows that ENG expression in tumor cells indirectly stimulates angiogenesis within the TME. Our results therefore support a dual role for tumor cell-specific ENG in MPNSTs (1) promoting proliferation and metastasis of tumor cells and (2) favoring the secretion of pro-angiogenic factors.

Notably, this study demonstrates that ENG may be a promising therapeutic target for MPNSTs irrespective of NF1 status, since targeting tumor and endothelial ENG with TRC105/M1043 inhibited tumor growth and metastasis in both NF1-associated (ST88-14) and sporadic (STS26T) MPNST xenograft models. Pharmacological ENG inhibition resulted in reduced *in vivo* proliferation and metastatic ability of MPNST cells and impaired tumor angiogenesis, which is consistent with our data from the *ENG* knockdown model and, therefore, supports the involvement of ENG in these biological processes in MPNSTs. Since, in addition to tumor cells and ECs, ENG expression has been reported in other cell types within the TME (e.g. fibroblasts) (43), we cannot rule out the possibility that the therapeutic effects of anti-ENG therapy in our MPNST xenografts are also due to ENG inhibition in these stromal cells. In line with our findings, several studies revealed that anti-ENG mAbs efficiently suppress tumor growth and metastasis in different mouse tumor models (18). Indeed, in phase I-III trials, TRC105, alone or in combination therapy, has shown a favorable safety profile and promising efficacy in patients with diverse tumor types, mainly in those with some advanced STS subtypes (18,44,45). Here, our preclinical data support a novel use of TRC105 for the treatment of MPNSTs.

Single-agent therapies have so far proved limited clinical success in MPNSTs (6,8), highlighting the importance of identifying successful combinatorial treatments to increase efficacy and prevent resistance. MEKis represent the most effective targeted therapy for patients with PNs, which may evolve to MPNSTs (19). Accordingly, MEK inhibition was recently shown to achieve complete response in a patient with recurrent and metastatic MPNST (46). However, extensive *in vivo* evidence has indicated robust yet transient anti-tumor effects of MEK inhibition alone in MPNSTs, suggesting that MEKis may prove to be more efficacious in combination with other agents (8). Our results demonstrate that the combination of the anti-ENG mAbs TRC105/M1043 with PD-0325901 exerts better therapeutic efficacy than MEKi monotherapy, resulting in sustained inhibition of MPNST growth. We found that ENG targeting cooperated with MEK inhibition to block ENG-Smad1/5 and MAPK/ERK pathway activation in MPNST cells and to reduce tumor cell proliferation and angiogenesis in primary tumors. These data support the notion that cross-talk between ENG-Smad1/5 and MAPK/ERK pathways contributes to MPNST growth, providing evidence for combining anti-ENG and anti-MEK agents as a novel improved therapeutic strategy against MPNSTs.

The inhibitory effects of the dual therapy on both primary tumor growth and established metastases suggest that the treatment could be effective for localized or metastatic disease. These findings have important translational implications, since intervention timing and disease staging have been reported as possible causes of failure of clinical trials involving MPNST patients (20). Importantly, several phase I-II studies have demonstrated the feasibility, safety and preliminary efficacy of combining anti-angiogenic agents (e.g. sorafenib or pazopanib) with a MEKi in patients with different solid malignancies, mainly in patients with other STS subtypes (8). TRC105 is emerging as a more promising strategy than these classic anti-angiogenic drugs for the treatment of cancer patients (12). Indeed, the clinical effects of TRC105 have been attributed to the inhibition of angiogenesis (47) but also to the targeting of tumor cells and immunosuppressive cells (e.g. T and B regulatory cells), which has been associated with reduced metastasis and better survival (48-50). These multi-targets effects of TRC105 contribute to its favorable efficacy in cancer patients (18). These data, together with our *in vivo* results, provide support for clinical investigation of the combination of TRC105 with MEKi as a novel therapy for MPNSTs.

In summary, our findings unveil a tumor-promoting function of ENG in MPNSTs and support the use of this protein as a novel biomarker and a promising therapeutic target for this disease. Notably, we also provide evidence that dual targeting of ENG and MEK may represent an attractive treatment option for MPNST patients.

## Acknowledgments

We apologize to those authors whose work could not be cited due to size limitations. We thank to Dr. David Lyden and Dr. Eduard Serra for their support in the project, we also thank Héctor Tejero for his help in analyzing RNAseq data. Dr. Peinado laboratory is funded by US Department of Defense (W81XWH-16-1-0131), Agencia Estatal de Investigación/Ministerio de Ciencia e Innovación (AEI/MCIN) (PID2020-118558RB-I00/AEI/10.13039/501100011033), Fundación Proyecto Neurofibromatosis and Fundación Científica AECC. We are also grateful for the support of the Ministerio de Universidades (Programa de Formación de Profesorado Universitario (FPU)) for the fellowship FPU016/05356 awarded to T.G-M and to the Translational NeTwork for the CLinical application of Extracellular VesicleS (TeNTaCLES) RED2018-102411-T(AEI/10.13039/501100011033). Angela Di Giannatale was supported during this work by a research grant from Nuovo-Soldati Foundation. The CNIO, certified as Severo Ochoa Excellence Centre, is supported by the Spanish Government through the Instituto de Salud Carlos III (ISCIII).

## Figure legends

**Supplementary figure 1.**
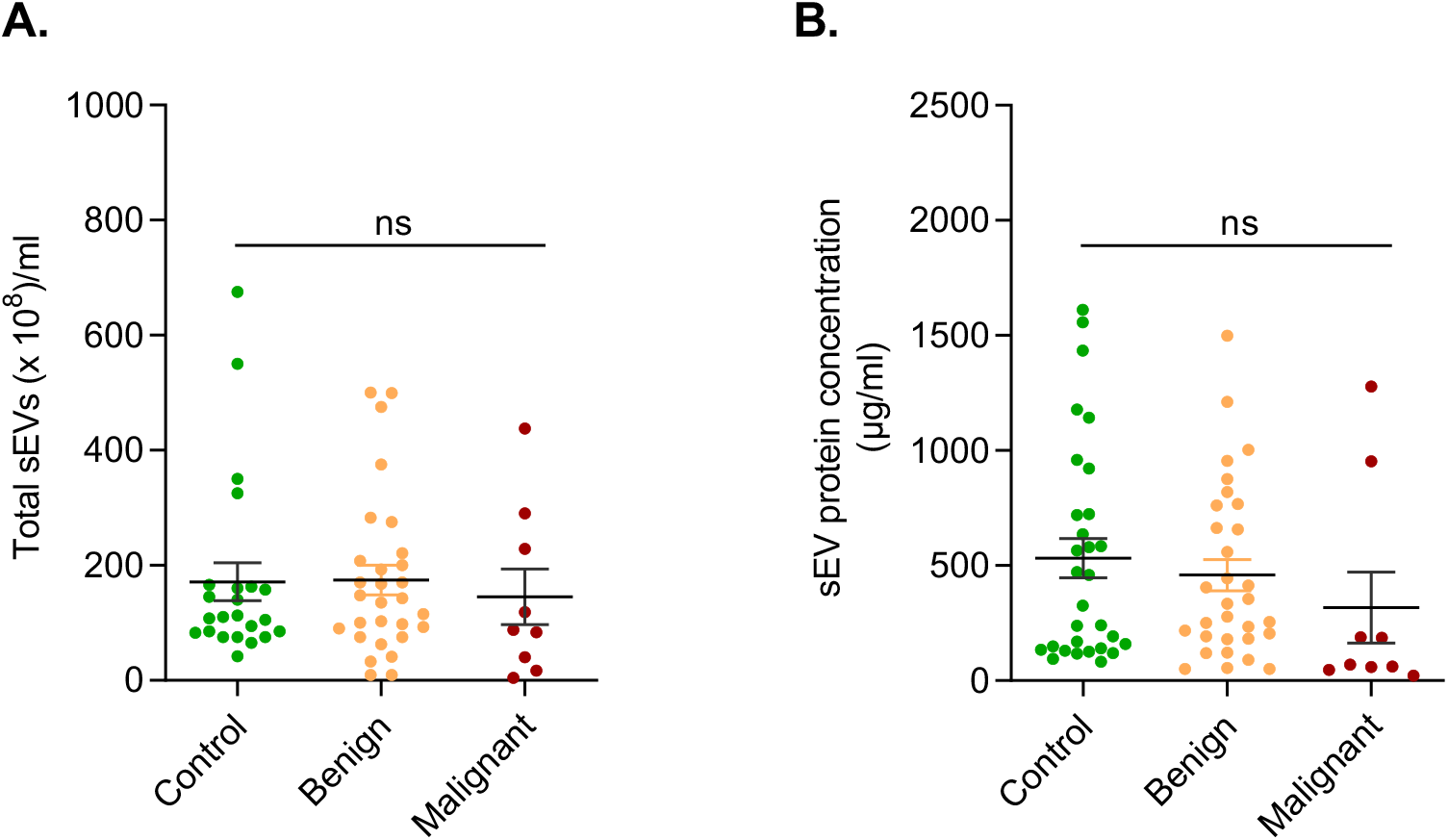
Characterization of human circulating sEVs. **A)** Quantification of total number of particles per milliliter in plasma samples using Nanoparticle Tracking Analysis (Nanosight). **B)** Protein concentration in circulating sEVs collected from the plasma of healthy controls (n=23) and patients with benign (PNs, n=33) or malignant (MPNSTs, n=11) PNSTs. Mean ± s.e.m.; ns, not significant; ns, not significant by one-way ANOVA.

**Supplementary figure 2.**
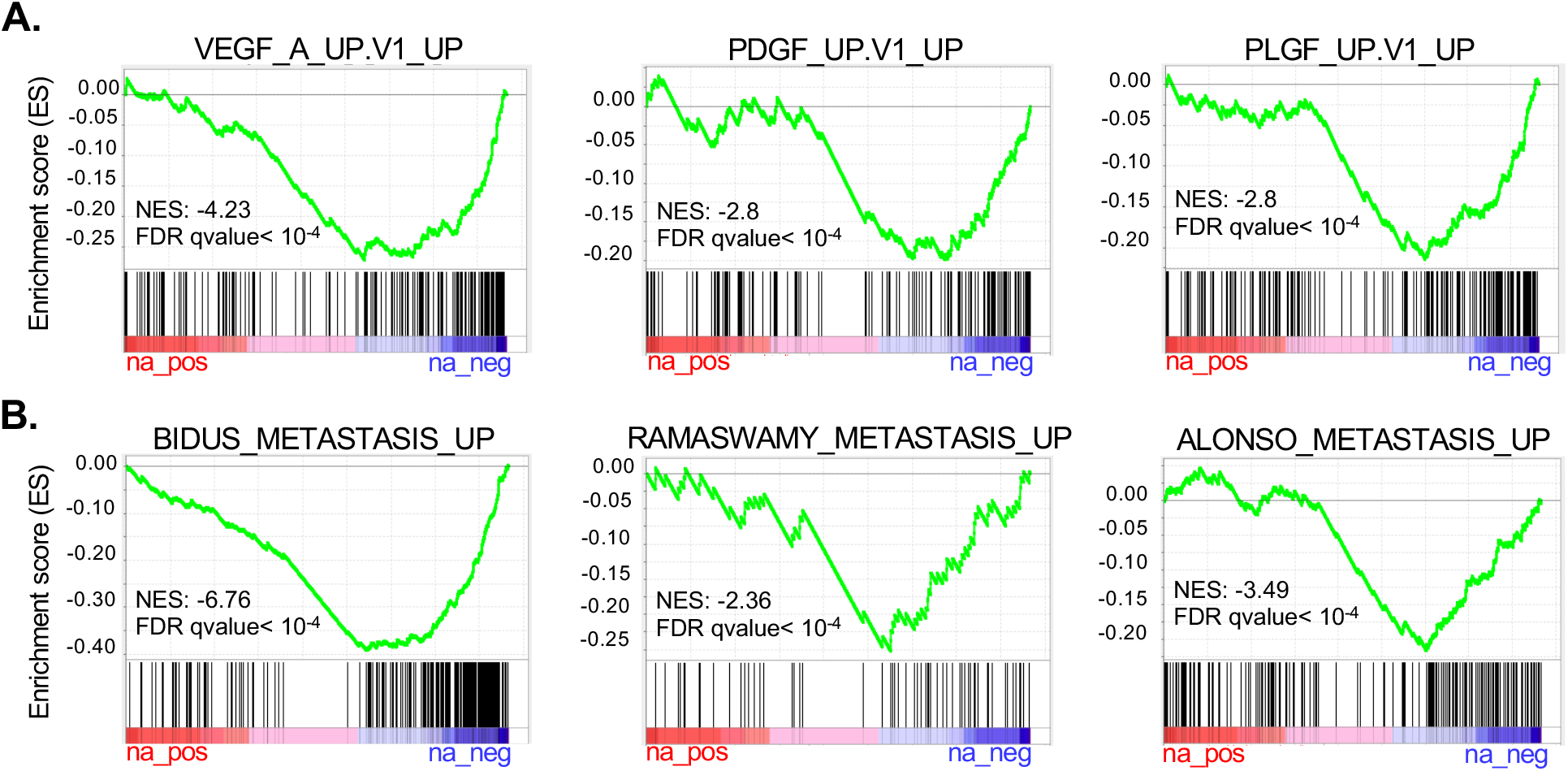
*ENG* knockdown leads to downregulation of pro-angiogenic and pro-metastatic gene signatures in MPNST cells. **A-B)** GSEA plots for the indicated gene signatures associated with pro-angiogenic factors (**A**) and metastasis (**B**) in shENG STS26T cells compared to shScramble cells. Signatures were obtained from Molecular Signatures Database (MSigDB).

**Supplementary figure 3.**
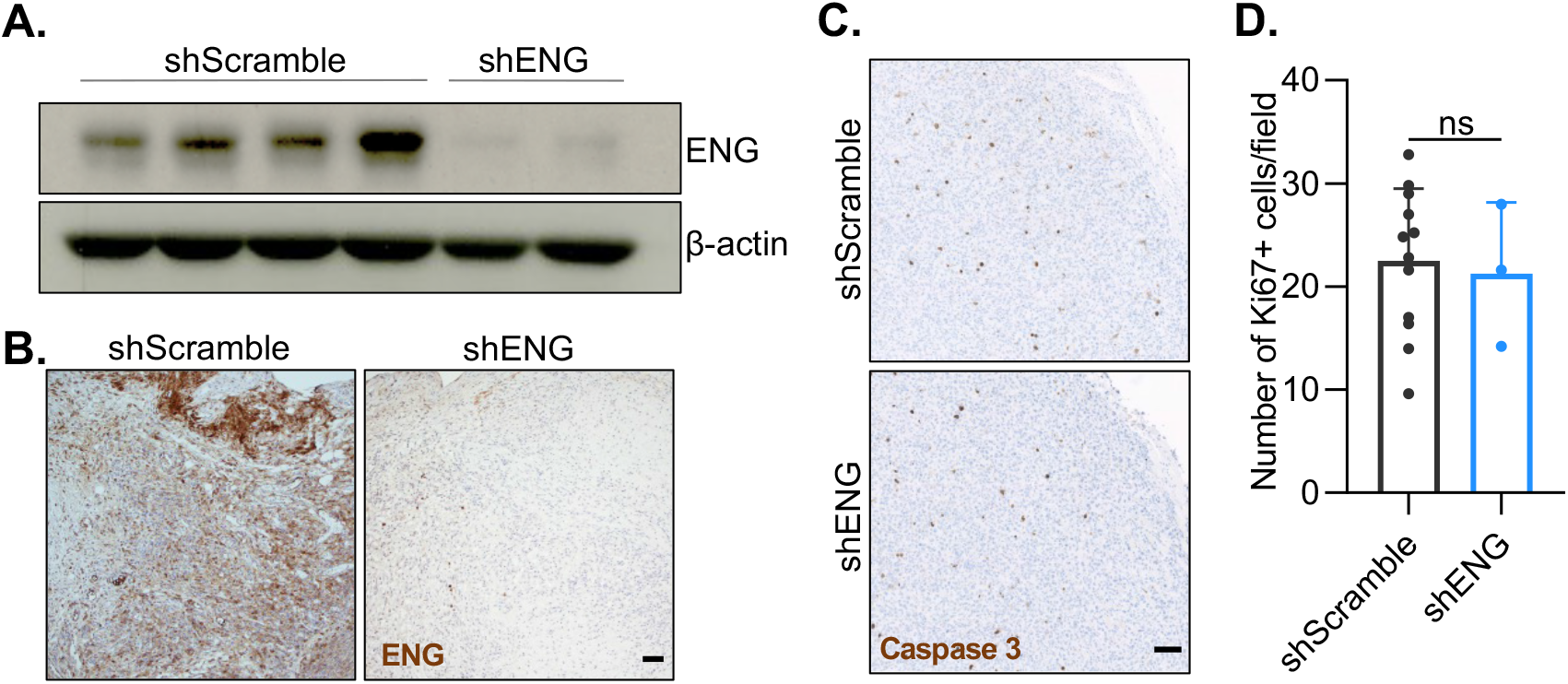
*ENG* knockdown does not affect MPNST cell apoptosis *in vivo*. **A)** Western blot analysis of ENG in shScramble- and shENG-STS26T xenografts. **B)** Representative images of ENG immunohistochemical staining in xenografts at end-point (day 32). Scale bar=200μm. **C, D)** IHC analysis of the apoptotic marker active-caspase 3 in the indicated STS26T xenografts (Scale bar=100μm) (**C**), and quantification (**D**). Dots in the graphs represent the mean number of positive cells per field of each tumor (5 fields/tumor). Mean ± s.e.m.; ns, not significant by unpaired t-test.

**Supplementary figure 4.**
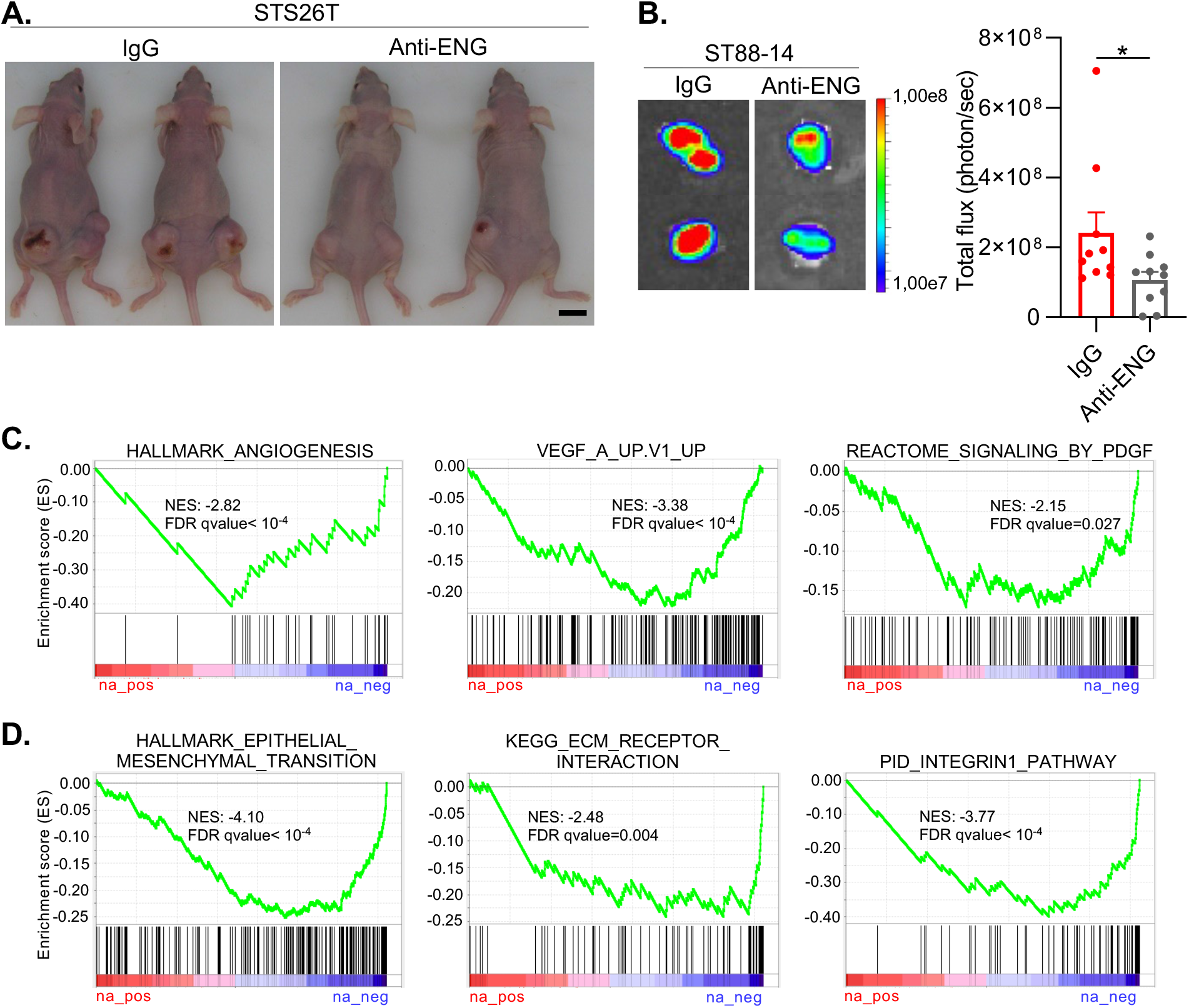
Anti-ENG therapy is active against MPNST xenografts. **A)** Representative images of mice bearing STS26T xenografts treated with IgG or TRC105/M1043 at day 28 (end point). Scale bar=1cm. n= 5 mice per group. **B)** Representative bioluminescence images (left panels) and total flux quantification (right panel) in control- and TRC105/M1043-treated ST88-14 tumors at day 28 (end point). Data correspond to mean ± s.e.m. of 10 tumors per condition; * P<0.05 by unpaired t-test. **C-D)** GSEA plots for the indicated gene sets related to angiogenesis (**C**) and metastasis (**D**) available in MSigDB (Anti-ENG versus IgG).

**Supplementary figure 5.**
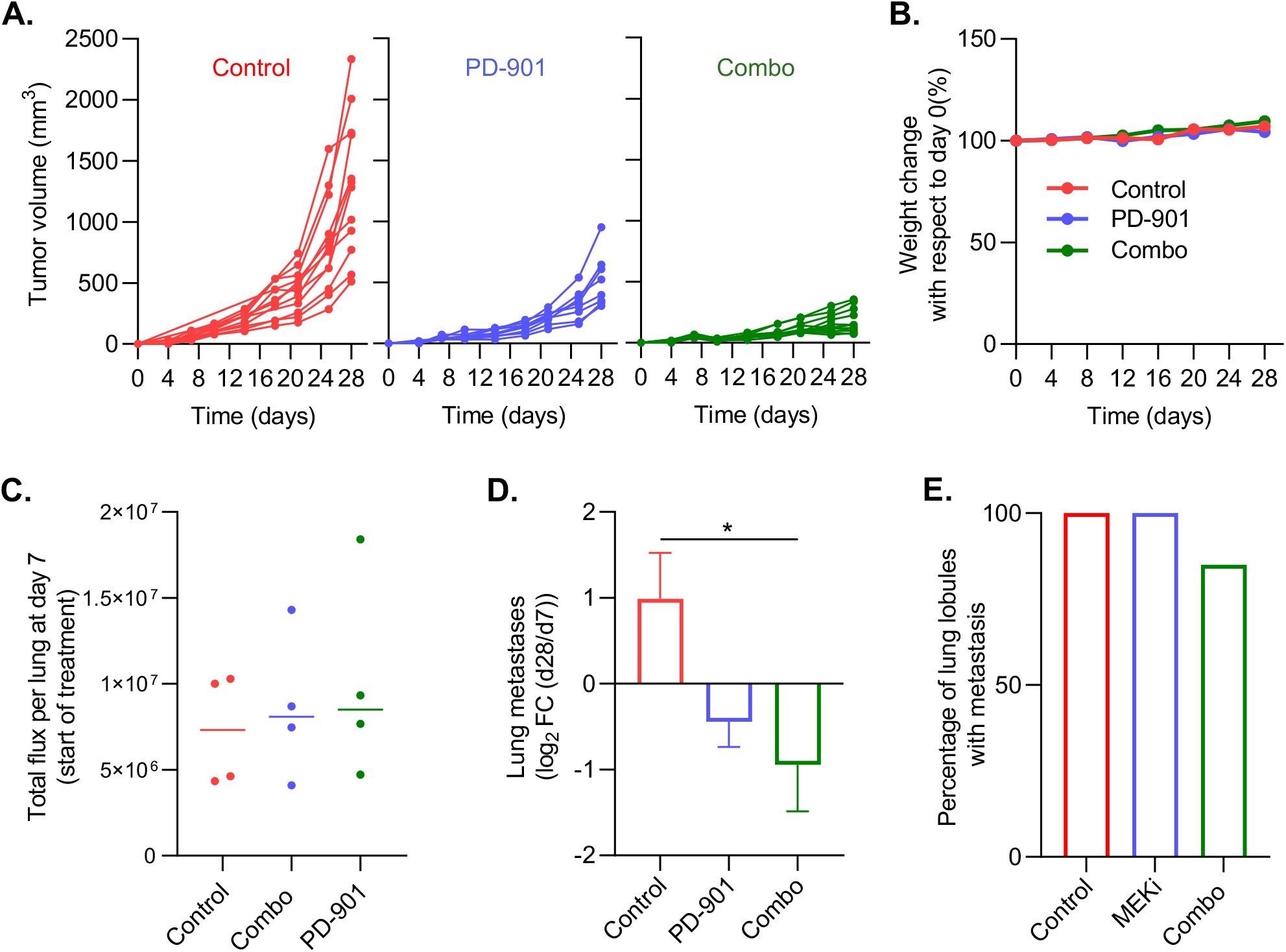
Combined anti-ENG/anti-MEK therapy inhibits MPNST growth without toxicity *in vivo*. **A)** Plots showing growth of individual subcutaneous STS26T tumors in mice treated with IgG plus vehicle (control), PD-0325901 (PD-901) alone or the combination of PD-901 with TRC105/M1043 (combo). **B)** Percentage of body weight changes relative to day 0 along the treatment period. **C)** Randomization in the lung metastasis experiment based on lung bioluminescent signal before start of treatment (day 7 post-tumor cell injection). **D)** Changes in lung metastatic burden quantified by luciferin photon flux at 21 days after the indicated treatments. Data are presented as log2 fold change of total flux at day 28 (end-point) compared to day 7 (start of treatment). Mean ± s.e.m. of four mice per group; * P<0.05 by one-way ANOVA. **E)** Percentage of lung lobules with metastasis (bioluminiscent signal above background) in each group of treatment.

**Supplementary figure 6.**
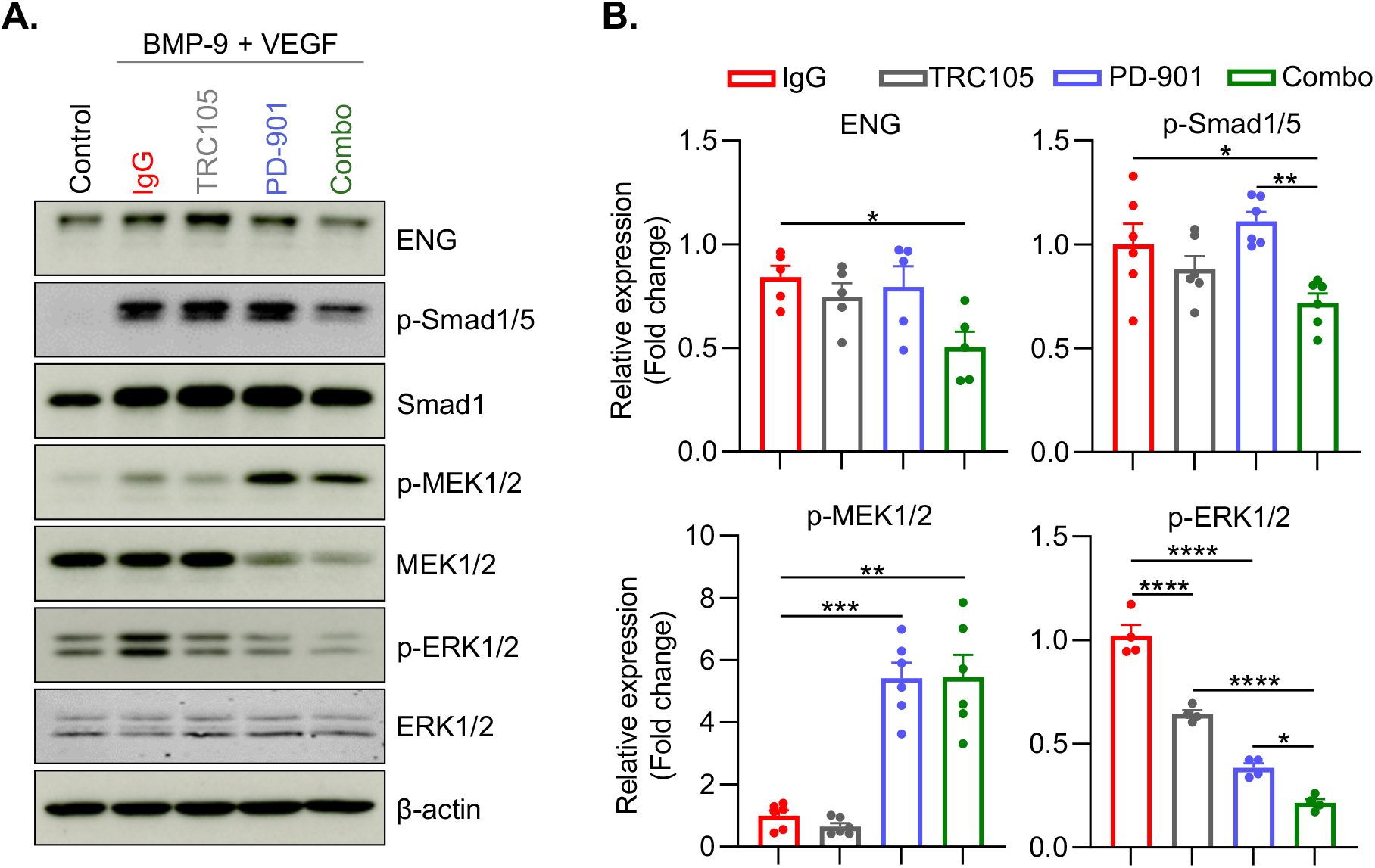
Dual inhibition of ENG and MEK efficiently blocks ENG-Smad1/5 and MAPK/ERK pathway activation in ST88-14 cells. **A, B)** Western blot analysis (**A**) and quantification (**B**) of ENG expression and Smad1/5, MEK and ERK phosphorylation in ST88-14 cells stimulated with BMP-9/VEGF in the presence of IgG control, the anti-human ENG mAb TRC105, the MEKi PD-0325901 (PD-901) or the combination of both drugs (combo). Cells without neither the pre-treatment nor BMP-9/VEGF stimulation were used as a control. Phosphorylated protein levels were normalized to the corresponding total protein levels. Data are presented as the fold change compared to IgG. Mean ± s.e.m. of at least four biological replicates; * P<0.05, ** P<0.01, *** P<0.001, **** P<0.0001 by one-way ANOVA

## ABBREVIATIONS

ANNUBP: Atypical neurofibromatosis neoplasms of uncertain biological potential
BCA: Bicinchoninic acid
BLI: Bioluminescence imaging
CBR: Clinical benefit rate
cfDNA: Circulating-free DNA
EC: Endothelial cell
ENG: Endoglin
ETP: Experimental Therapeutics Unit
EV: Extracellular vesicle
GAP: GTPase activating protein
GSEA: Gene set enrichment analysis
H&E: Hematoxylin and eosin
IHC: Immunohistochemistry
ISCIII: Instituto de Salud Carlos III
LN: Lymph node
MEKi: MEK inhibitor
MPNST: Malignant peripheral nerve sheath tumor
NF1: Neurofibromatosis type 1
NSG: NOD scid gamma
NTA: Nanoparticle tracking analysis
PD-901: PD-0325901
PN: Plexiform neurofibroma
PNST: Peripheral nerve sheath tumor
RNA-seq: RNA sequencing
sEV: Small extracellular vesicle
STS: Soft-tissue sarcoma
TMA: Tissue microarray
TME: Tumor microenvironment

## REFERENCES

1. Farid M, Demicco EG, Garcia R, Ahn L, Merola PR, Cioffi A, et al. Malignant peripheral nerve sheath tumors. Oncologist 2014;19(2):193–201 doi 10.1634/theoncologist.2013-0328.

2. Korfhage J, Lombard DB. Malignant Peripheral Nerve Sheath Tumors: From Epigenome to Bedside. Mol Cancer Res 2019;17(7):1417–28 doi 10.1158/1541-7786.MCR-19-0147.

3. Gutmann DH, Ferner RE, Listernick RH, Korf BR, Wolters PL, Johnson KJ. Neurofibromatosis type 1. Nat Rev Dis Primers 2017;3:17004 doi 10.1038/nrdp.2017.4.

4. Prudner BC, Ball T, Rathore R, Hirbe AC. Diagnosis and management of malignant peripheral nerve sheath tumors: Current practice and future perspectives. Neurooncol Adv 2020;2(Suppl 1):i40–i9 doi 10.1093/noajnl/vdz047.

5. Bradford D, Kim A. Current treatment options for malignant peripheral nerve sheath tumors. Curr Treat Options Oncol 2015;16(3):328 doi 10.1007/s11864-015-0328-6.

6. Martin E, Lamba N, Flucke UE, Verhoef C, Coert JH, Versleijen-Jonkers YMH, et al. Non-cytotoxic systemic treatment in malignant peripheral nerve sheath tumors (MPNST): A systematic review from bench to bedside. Crit Rev Oncol Hematol 2019;138:223–32 doi 10.1016/j.critrevonc.2019.04.007.

7. Mohamad T, Plante C, Brosseau JP. Toward Understanding the Mechanisms of Malignant Peripheral Nerve Sheath Tumor Development. Int J Mol Sci 2021;22(16) doi 10.3390/ijms22168620.

8. Gonzalez-Munoz T, Kim A, Ratner N, Peinado H. The need for new treatments targeting MPNST: the potential of strategies combining MEK inhibitors with antiangiogenic agents. Clin Cancer Res 2022 doi 10.1158/1078-0432.CCR-21-3760.

9. Wasa J, Nishida Y, Suzuki Y, Tsukushi S, Shido Y, Hosono K, et al. Differential expression of angiogenic factors in peripheral nerve sheath tumors. Clin Exp Metastasis 2008;25(7):819–25 doi 10.1007/s10585-008-9197-8.

10. Nishida Y, Urakawa H, Nakayama R, Kobayashi E, Ozaki T, Ae K, et al. Phase II clinical trial of pazopanib for patients with unresectable or metastatic malignant peripheral nerve sheath tumors. Int J Cancer 2021;148(1):140–9 doi 10.1002/ijc.33201.

11. Lebrin F, Goumans MJ, Jonker L, Carvalho RL, Valdimarsdottir G, Thorikay M, et al. Endoglin promotes endothelial cell proliferation and TGF-beta/ALK1 signal transduction. EMBO J 2004;23(20):4018–28 doi 10.1038/sj.emboj.7600386.

12. Gonzalez Munoz T, Amaral AT, Puerto-Camacho P, Peinado H, de Alava E. Endoglin in the Spotlight to Treat Cancer. Int J Mol Sci 2021;22(6) doi 10.3390/ijms22063186.

13. Rosen LS, Gordon MS, Robert F, Matei DE. Endoglin for targeted cancer treatment. Curr Oncol Rep 2014;16(2):365 doi 10.1007/s11912-013-0365-x.

14. Graña O, Rubio-Camarillo M, Fdez-Riverola F, Pisano DG, Glez-Peña D. Nextpresso: Next Generation Sequencing Expression Analysis Pipeline. Current Bioinformatics 2018;13(6):583–91 doi http://dx.doi.org/10.2174/1574893612666170810153850.

15. Dahlberg WK, Little JB, Fletcher JA, Suit HD, Okunieff P. Radiosensitivity in vitro of human soft tissue sarcoma cell lines and skin fibroblasts derived from the same patients. Int J Radiat Biol 1993;63(2):191–8 doi 10.1080/09553009314550251.

16. Glover TW, Stein CK, Legius E, Andersen LB, Brereton A, Johnson S. Molecular and cytogenetic analysis of tumors in von Recklinghausen neurofibromatosis. Genes Chromosomes Cancer 1991;3(1):62–70 doi 10.1002/gcc.2870030111.

17. Lee NY, Blobe GC. The interaction of endoglin with beta-arrestin2 regulates transforming growth factor-beta-mediated ERK activation and migration in endothelial cells. J Biol Chem 2007;282(29):21507–17 doi 10.1074/jbc.M700176200.

18. Liu Y, Paauwe M, Nixon AB, Hawinkels L. Endoglin Targeting: Lessons Learned and Questions That Remain. Int J Mol Sci 2020;22(1) doi 10.3390/ijms22010147.

19. Casey D, Demko S, Sinha A, Mishra-Kalyani PS, Shen YL, Khasar S, et al. FDA Approval Summary: Selumetinib for Plexiform Neurofibroma. Clin Cancer Res 2021;27(15):4142–6 doi 10.1158/1078-0432.CCR-20-5032.

20. Kim A, Stewart DR, Reilly KM, Viskochil D, Miettinen MM, Widemann BC. Malignant Peripheral Nerve Sheath Tumors State of the Science: Leveraging Clinical and Biological Insights into Effective Therapies. Sarcoma 2017;2017:7429697 doi 10.1155/2017/7429697.

21. Ghadimi MP, Young ED, Belousov R, Zhang Y, Lopez G, Lusby K, et al. Survivin is a viable target for the treatment of malignant peripheral nerve sheath tumors. Clin Cancer Res 2012;18(9):2545–57 doi 10.1158/1078-0432.CCR-11-2592.

22. Nolan-Stevaux O, Zhong W, Culp S, Shaffer K, Hoover J, Wickramasinghe D, et al. Endoglin requirement for BMP9 signaling in endothelial cells reveals new mechanism of action for selective anti-endoglin antibodies. PLoS One 2012;7(12):e50920 doi 10.1371/journal.pone.0050920.

23. Dhillon AS, Hagan S, Rath O, Kolch W. MAP kinase signalling pathways in cancer. Oncogene 2007;26(22):3279–90 doi 10.1038/sj.onc.1210421.

24. Wang J, Pollard K, Allen AN, Tomar T, Pijnenburg D, Yao Z, et al. Combined Inhibition of SHP2 and MEK Is Effective in Models of NF1-Deficient Malignant Peripheral Nerve Sheath Tumors. Cancer Res 2020;80(23):5367–79 doi 10.1158/0008-5472.CAN-20-1365.

25. Sakamoto R, Kajihara I, Miyauchi H, Maeda-Otsuka S, Yamada-Kanazawa S, Sawamura S, et al. Inhibition of Endoglin Exerts Antitumor Effects through the Regulation of Non-Smad TGF-beta Signaling in Angiosarcoma. J Invest Dermatol 2020;140(10):2060–72 e6 doi 10.1016/j.jid.2020.01.031.

26. Mitsui H, Shibata K, Mano Y, Suzuki S, Umezu T, Mizuno M, et al. The expression and characterization of endoglin in uterine leiomyosarcoma. Clin Exp Metastasis 2013;30(6):731–40 doi 10.1007/s10585-013-9574-9.

27. Boeuf S, Bovee JV, Lehner B, van den Akker B, van Ruler M, Cleton-Jansen AM, et al. BMP and TGFbeta pathways in human central chondrosarcoma: enhanced endoglin and Smad 1 signaling in high grade tumors. BMC Cancer 2012;12:488 doi 10.1186/1471-2407-12-488.

28. Gromova P, Rubin BP, Thys A, Cullus P, Erneux C, Vanderwinden JM. ENDOGLIN/CD105 is expressed in KIT positive cells in the gut and in gastrointestinal stromal tumours. J Cell Mol Med 2012;16(2):306–17 doi 10.1111/j.1582-4934.2011.01315.x.

29. Pardali E, van der Schaft DW, Wiercinska E, Gorter A, Hogendoorn PC, Griffioen AW, et al. Critical role of endoglin in tumor cell plasticity of Ewing sarcoma and melanoma. Oncogene 2011;30(3):334–45 doi 10.1038/onc.2010.418.

30. Cescon DW, Bratman SV, Chan SM, Siu LL. Circulating tumor DNA and liquid biopsy in oncology. Nat Cancer 2020;1(3):276–90 doi 10.1038/s43018-020-0043-5.

31. Szymanski JJ, Sundby RT, Jones PA, Srihari D, Earland N, Harris PK, et al. Cell-free DNA ultra-low-pass whole genome sequencing to distinguish malignant peripheral nerve sheath tumor (MPNST) from its benign precursor lesion: A cross-sectional study. PLoS Med 2021;18(8):e1003734 doi 10.1371/journal.pmed.1003734.

32. Li C, Guo B, Wilson PB, Stewart A, Byrne G, Bundred N, et al. Plasma levels of soluble CD105 correlate with metastasis in patients with breast cancer. Int J Cancer 2000;89(2):122–6 doi 10.1002/(sici)1097-0215(20000320)89:2<122::aid-ijc4>3.0.co;2-m.

33. Svatek RS, Karam JA, Roehrborn CG, Karakiewicz PI, Slawin KM, Shariat SF. Preoperative plasma endoglin levels predict biochemical progression after radical prostatectomy. Clin Cancer Res 2008;14(11):3362–6 doi 10.1158/1078-0432.CCR-07-4707.

34. Nogues A, Gallardo-Vara E, Zafra MP, Mate P, Marijuan JL, Alonso A, et al. Endoglin (CD105) and VEGF as potential angiogenic and dissemination markers for colorectal cancer. World J Surg Oncol 2020;18(1):99 doi 10.1186/s12957-020-01871-2.

35. Douglas SR, Yeung KT, Yang J, Blair SL, Cohen O, Eliceiri BP. Identification of CD105+ Extracellular Vesicles as a Candidate Biomarker for Metastatic Breast Cancer. J Surg Res 2021;268:168–73 doi 10.1016/j.jss.2021.06.050.

36. Bernabeu C, Lopez-Novoa JM, Quintanilla M. The emerging role of TGF-beta superfamily coreceptors in cancer. Biochim Biophys Acta 2009;1792(10):954–73 doi 10.1016/j.bbadis.2009.07.003.

37. Santibanez JF, Perez-Gomez E, Fernandez LA, Garrido-Martin EM, Carnero A, Malumbres M, et al. The TGF-beta co-receptor endoglin modulates the expression and transforming potential of H-Ras. Carcinogenesis 2010;31(12):2145–54 doi 10.1093/carcin/bgq199.

38. Ahsan S, Ge Y, Tainsky MA. Combinatorial therapeutic targeting of BMP2 and MEK-ERK pathways in NF1-associated malignant peripheral nerve sheath tumors. Oncotarget 2016;7(35):57171–85 doi 10.18632/oncotarget.11036.

39. Postiglione L, Di Domenico G, Caraglia M, Marra M, Giuberti G, Del Vecchio L, et al. Differential expression and cytoplasm/membrane distribution of endoglin (CD105) in human tumour cell lines: Implications in the modulation of cell proliferation. Int J Oncol 2005;26(5):1193–201 doi 10.3892/ijo.26.5.1193.

40. Paauwe M, Heijkants RC, Oudt CH, van Pelt GW, Cui C, Theuer CP, et al. Endoglin targeting inhibits tumor angiogenesis and metastatic spread in breast cancer. Oncogene 2016;35(31):4069–79 doi 10.1038/onc.2015.509.

41. Romero D, O’Neill C, Terzic A, Contois L, Young K, Conley BA, et al. Endoglin regulates cancer-stromal cell interactions in prostate tumors. Cancer Res 2011;71(10):3482–93 doi 10.1158/0008-5472.CAN-10-2665.

42. Duwel A, Eleno N, Jerkic M, Arevalo M, Bolanos JP, Bernabeu C, et al. Reduced tumor growth and angiogenesis in endoglin-haploinsufficient mice. Tumour Biol 2007;28(1):1–8 doi 10.1159/000097040.

43. Paauwe M, Schoonderwoerd MJA, Helderman R, Harryvan TJ, Groenewoud A, van Pelt GW, et al. Endoglin Expression on Cancer-Associated Fibroblasts Regulates Invasion and Stimulates Colorectal Cancer Metastasis. Clin Cancer Res 2018;24(24):6331–44 doi 10.1158/1078-0432.CCR-18-0329.

44. Attia S, Sankhala KK, Riedel RF, Robinson SI, Conry RM, Boland PM, et al. A phase 1B/ phase 2A study of TRC105 (Endoglin Antibody) in combination with pazopanib (P) in patients (pts) with advanced soft tissue sarcoma (STS). Journal of Clinical Oncology 2016;34(15_suppl):11016- doi 10.1200/JCO.2016.34.15_suppl.11016.

45. Mehta CR, Liu L, Theuer C. An adaptive population enrichment phase III trial of TRC105 and pazopanib versus pazopanib alone in patients with advanced angiosarcoma (TAPPAS trial). Ann Oncol 2019;30(1):103–8 doi 10.1093/annonc/mdy464.

46. Nagabushan S, Lau LMS, Barahona P, Wong M, Sherstyuk A, Marshall GM, et al. Efficacy of MEK inhibition in a recurrent malignant peripheral nerve sheath tumor. NPJ Precis Oncol 2021;5(1):9 doi 10.1038/s41698-021-00145-8.

47. Liu Y, Starr MD, Brady JC, Dellinger A, Pang H, Adams B, et al. Modulation of circulating protein biomarkers following TRC105 (anti-endoglin antibody) treatment in patients with advanced cancer. Cancer Med 2014;3(3):580–91 doi 10.1002/cam4.207.

48. Rosen LS, Hurwitz HI, Wong MK, Goldman J, Mendelson DS, Figg WD, et al. A phase I first-in-human study of TRC105 (Anti-Endoglin Antibody) in patients with advanced cancer. Clin Cancer Res 2012;18(17):4820–9 doi 10.1158/1078-0432.CCR-12-0098.

49. Apolo AB, Karzai FH, Trepel JB, Alarcon S, Lee S, Lee MJ, et al. A Phase II Clinical Trial of TRC105 (Anti-Endoglin Antibody) in Adults With Advanced/Metastatic Urothelial Carcinoma. Clin Genitourin Cancer 2017;15(1):77–85 doi 10.1016/j.clgc.2016.05.010.

50. Karzai FH, Apolo AB, Cao L, Madan RA, Adelberg DE, Parnes H, et al. A phase I study of TRC105 anti-endoglin (CD105) antibody in metastatic castration-resistant prostate cancer. BJU Int 2015;116(4):546–55 doi 10.1111/bju.12986.

